# Spatiotemporal Control of Pathway Sensors and Cross-Pathway Feedback Regulate a Cell Differentiation MAPK Pathway in Yeast

**DOI:** 10.1101/2020.12.18.423530

**Authors:** Aditi Prabhakar, Beatriz Gonzalez, Heather Dionne, Sukanya Basu, Paul J. Cullen

**Affiliations:** Department of Biological Sciences, University at Buffalo, Buffalo, NY 14260-1300

**Author notes:** Corresponding author: Paul J. Cullen, Address: Department of Biological Sciences, 532 Cooke Hall, State University of New York at Buffalo, Buffalo, NY 14260-1300. Author contributions: A.P. designed experiments, generated data, and wrote the paper; B.G. generated data; S.B. generated data; H.D. designed experiments; P.J.C. designed experiments and wrote the paper. The authors have no competing interests in the study.

**Keywords:** mucins, cell cycle, mother-bud neck, septins, cell synchronization, HOG, filamentous growth, invasive growth, pseudohyphal growth

## Abstract

Mitogen-Activated Protein Kinase (MAPK) pathways control cell differentiation and the response to stress. MAPK pathways can share components with other pathways yet induce specific responses through mechanisms that remain unclear. In *Saccharomyces cerevisiae*, the MAPK pathway that controls filamentous growth (fMAPK) shares components with the MAPK pathway that regulates the response to osmotic stress (HOG). By exploring temporal regulation of MAPK signaling, we show here that the two pathways exhibited different patterns of activity throughout the cell cycle. The different patterns resulted from different expression profiles of genes encoding the mucin sensors (*MSB2* for fMAPK and *HKR1* for HOG). We also show that positive feedback through the fMAPK pathway stimulated the HOG pathway, presumably to modulate fMAPK pathway activity. By exploring spatial regulation of MAPK signaling, we found that the shared tetraspan protein, Sho1p, which has a dynamic localization pattern, induced the fMAPK pathway at the mother-bud neck. A Sho1p-interacting protein, Hof1p, which also localizes to the mother-bud neck and regulates cytokinesis, also regulated the fMAPK pathway. Therefore, spatial and temporal regulation of pathway sensors, and cross-pathway feedback, regulate a MAPK pathway that controls a cell differentiation response in yeast.

## INTRODUCTION

Mitogen activated protein (MAP) kinase pathways are evolutionary conserved signaling modules that control growth (Johnson and Lapadat 2002; Lavoie *et al.* 2020), cell differentiation (Chen and Thorner 2007; Raman *et al.* 2007; Dinsmore and Soriano 2018), and the response to stress (Roux and Blenis 2004; Yoon and Seger 2006; Shaul and Seger 2007; Papa *et al.* 2019). Like other signaling pathways, MAP kinase pathways operate in interconnected networks where inputs from multiple pathways converge (Jordan *et al.* 2000; Fischer *et al.* 2018). Moreover, MAPK pathways are subject to exquisite spatiotemporal control, in that signaling occurs at a precise place, typically on the cell surface, and for a defined duration to elicit an appropriate response. Mis-regulation of MAPK pathways is linked to various diseases such as cancers, polycystic kidney disease, obesity, diabetes and developmental disorders (Lee *et al.* 2000; Hirosumi *et al.* 2002; Maekawa *et al.* 2005; Omori *et al.* 2006; Rodriguez-Viciana *et al.* 2006; Roberts and Der 2007; Lawrence *et al.* 2008). Therefore, understanding how MAPK pathways induce a specific signal an integrated network remains an important open question.

In the budding yeast *Saccharomyces cerevisiae*, MAP kinase pathways control well-characterized cell differentiation responses that occur in response to extrinsic cues (e.g mating and filamentous growth). Yeast MAP kinase pathways also control the response to stresses like high osmolarity (HOG). In response to nutrient limitation, yeast and other fungal species can undergo filamentous (or invasive/pseudohyphal) growth, where cells differentiate into interconnected and elongated filaments (Gimeno *et al.* 1992; Roberts and Fink 1994; Mosch *et al.* 1996; Peter *et al.* 1996; Leberer *et al.* 1997; Pan *et al.* 2000; Gancedo 2001; Adhikari *et al.* 2015b). In some fungal pathogens, this microbial differentiation response is critical for virulence (Lo *et al.* 1997). In yeast, the MAP kinase pathway that regulates filamentous growth (**Fig. 1A**, fMAPK, green) is controlled by the mucin-type glycoprotein Msb2p (Cullen *et al.* 2004). The underglycosylation of Msb2p occurs under nutrient-limiting conditions, which results in proteolytic processing and release of its inhibitory extracellular domain by an aspartyl-type protease, Yps1p (Adhikari *et al.* 2015b). Msb2p functions with a four-pass (tetraspan) adaptor protein, Sho1p (O’Rourke and Herskowitz 1998; Cullen *et al.* 2004), and another adaptor, Opy2p (Yamamoto *et al.* 2010; Karunanithi and Cullen 2012). Msb2p and Sho1p converge on the Rho GTPase Cdc42p, which regulates the fMAPK pathway by binding to and activating the p21-activated (PAK) kinase, Ste20p (Peter *et al.* 1996; Leberer *et al.* 1997; Johnson 1999; Bi and Park 2012).

**Figure 1.**
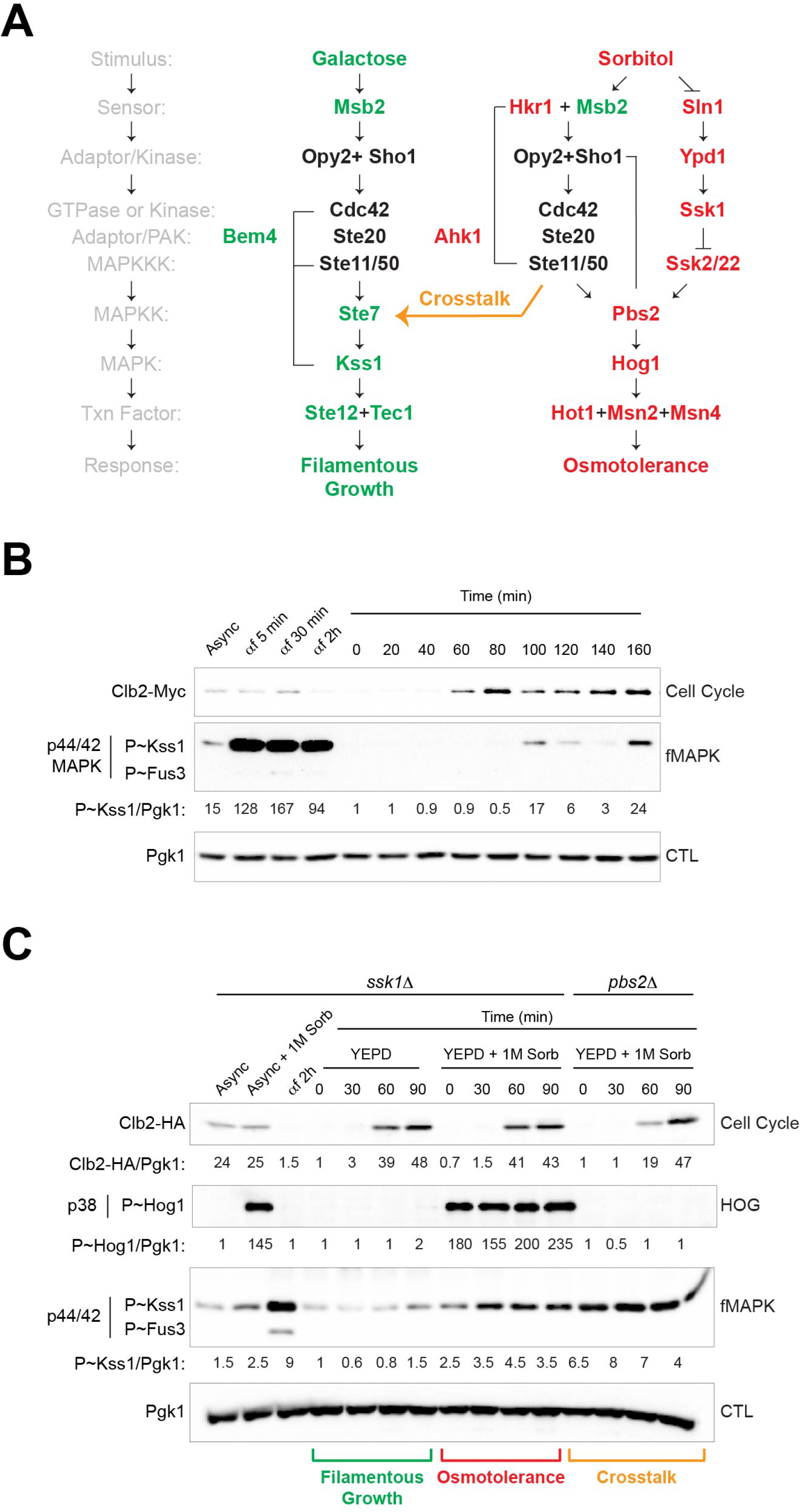
Two Cdc42p-dependent pathways in yeast share components yet induce different responses. (**A**) MAPK pathways that regulate filamentous growth (green, fMAPK) and osmolarity (red, HOG). Proteins shown in black (Opy2, Sho1, Cdc42, Ste20, Ste50, and Ste11) regulate both pathways. Black lines indicate interactions between adaptors and core pathway regulators. Orange arrow shows crosstalk between HOG and the fMAPK pathway, which occurs in cells lacking Pbs2p or Hog1p. (**B**) Immunoblot analysis of wild-type cells (PC7602) synchronized in G1 by α factor and released into YEPD medium. Samples were also harvested before (Async) and during α-factor treatment at indicated times (αf 5 min, 30 min and 2 h). Cell extracts were probed with antibodies to Clb2p-Myc (to monitor the cell cycle), P~Kss1p (p44/42, to monitor fMAPK), and Pgk1p as a control for protein levels (CTL). Numbers refer to the ratio of P~Kss1p to Pgk1p levels relative to time 0, which was set to 1. (**C**) Immunoblot analysis of *ssk1*Δ (PC7494) and *pbs2*Δ (PC7493) mutants synchronized in G1 by α factor. For the *ssk1*Δ mutant, synchronized cells were released into YEPD medium, and samples were harvested at the indicated times to measure basal HOG levels. Another aliquot was harvested and resuspended in YEPD supplemented with 1M sorbitol (YEPD + 1M Sorb) for 5 min before harvesting to measure activated HOG levels. Samples were also harvested before (Async) and during α-factor treatment (αf 2h). A sample of asynchronous culture was harvested and resuspended in YEPD supplemented with 1M sorbitol for 5 min (Async + 1M Sorb). For the *pbs2*Δ mutant, synchronized cells were released into YEPD medium. At indicated the times, aliquots were harvested and resuspended in YEPD supplemented with 1M sorbitol (YEPD + 1M Sorb) for 5 min before harvesting. Cell extracts were probed with antibodies to Clb2p-HA (to monitor the cell cycle), P~Kss1p (p44/42, to monitor fMAPK), P~Hog1p (p38, to monitor HOG), and Pgk1p, as a control for protein levels (CTL). Numbers refer to the ratio of Clb2p-HA to Pgk1p levels, P~Kss1p to Pgk1p levels and P~Hog1p to Pgk1p levels relative to time 0, which was set to 1.

When activated, Ste20p regulates a MAP kinase cascade composed of the MAPKKK (Ste11p), which phosphorylates and activates the MAPKK (Ste7p), which in turn phosphorylates the MAP kinase, Kss1p (Roberts and Fink 1994; Madhani *et al.* 1997). The active (phosphorylated) form of Kss1p regulates transcription factors including Ste12p and Tec1p, and the transcriptional repressor Dig1p, to induce target gene expression (Ma *et al.* 1995; Madhani and Fink 1997; Bardwell *et al.* 1998; Rupp *et al.* 1999; Roberts *et al.* 2000; van der Felden *et al.* 2014; Pelet 2017). The transcriptional targets of the fMAPK pathway encode products that induce differentiation to the filamentous cell type (Liu *et al.* 1993; Madhani and Fink 1997). Ste12p and Tec1p also stimulate the fMAPK pathway through positive feedback, because a subset of transcriptional targets of the fMAPK pathway encode pathway components (*MSB2, KSS1, STE12*, and *TEC1*)(Madhani *et al.* 1999; Roberts *et al.* 2000; Cullen *et al.* 2004; Adhikari and Cullen 2014).

The fMAPK pathway utilizes a subset of components that also function in other MAP kinase pathways, such as the HOG (O’Rourke and Herskowitz 2002; Cullen *et al.* 2004; Tatebayashi *et al.* 2006) and mating pathways (Bardwell *et al.* 1998; Sabbagh *et al.* 2001; Breitkreutz and Tyers 2002; Bardwell 2004; Schwartz and Madhani 2004; Bardwell 2006). The HOG pathway responds to changes in osmolarity and is composed of partially redundant branches (**Fig. 1A**). Like the fMAPK pathway, the Ste11p branch of the HOG pathway requires Sho1p, Cdc42p, Ste20p, and Ste11p (Tatebayashi *et al.* 2006). The pathways diverge after Ste11p, which phosphorylates Ste7p to regulate the fMAPK pathway, and Pbs2p to regulate the HOG pathway (Brewster *et al.* 1993; Maeda *et al.* 1994). Another difference between the pathways occurs at the level of the mucins, Msb2p and Hkr1p. Hkr1p regulates the HOG pathway, while Msb2p has been implicated in regulating both the fMAPK and HOG pathways [**Fig. 1A**; (O’rourke and Herskowitz 2002; Cullen *et al.* 2004; Tatebayashi *et al.* 2007; Pitoniak *et al.* 2009; Tanaka *et al.* 2014; Yamamoto *et al.* 2016)]. Msb2p and Hkr1p regulate the HOG pathway by different mechanisms (Tanaka *et al.* 2014). Importantly, Hkr1p does not regulate the fMAPK pathway, and overexpression of the mucins induce different target genes (Pitoniak *et al.* 2009). Moreover, the two pathways function in an antagonistic manner (Davenport *et al.* 1999; Adhikari and Cullen 2014). Therefore, selectivity in propagating a downstream signal presumably occurs at the level of the mucin glycoproteins and adaptors, although how this occurs is not well understood.

In this study, we explored the spatiotemporal regulation of the fMAPK and HOG pathways. One way that MAPK pathways exert precise temporal control is by regulating progression through the cell cycle. However, relatively little is known about how MAPK pathways are themselves regulated throughout the cell cycle. We previously found that the activity of the fMAPK pathway fluctuates throughout the cell cycle (Prabhakar *et al.* 2020), which provided an opportunity to explore this question in depth. We show here that the fMAPK and the HOG pathways show different patterns of cell-cycle regulation. While the activity of the fMAPK pathway peaked in M phase, the HOG pathway could be activated at any point in the cell cycle. The cell-cycle regulation of the fMAPK pathway stemmed from cell-cycle regulation of the mucin sensor, Msb2p. We also show that positive feedback through the fMAPK pathway induced the HOG pathway, generating cross-pathway feedback between the pathways. We have also previously shown that the fMAPK pathway is activated spatially by proteins that mark the cell poles (Basu *et al.* 2016). By examining the spatial localization of MAPK pathway sensors, we show that the tetraspan protein Sho1p was localized to the mother-bud neck during elevated fMAPK pathway activity. Sho1p-interacting proteins that also localize to the mother-bud neck were required for fMAPK pathway activity. Taken together, these results define new aspects of spatiotemporal regulation that leads to precise regulation of a MAPK pathway. Our findings may extend to other systems, where precise regulation of signaling pathways is required to induce cell differentiation.

## RESULTS

### MAPK Pathways that Share Components Show Different Patterns of Activity Throughout the Cell Cycle

Given that cell-cycle regulation of MAP kinase pathways is an under-explored aspect of MAP kinase pathway regulation, we explored the cell-cycle regulation of the fMAPK pathway that controls filamentous growth in yeast. In one approach, cells were synchronized by α-factor, which arrests cells in the G1 phase of the cell cycle (Elion *et al.* 1993; Peter *et al.* 1993; Peter and Herskowitz 1994). Release of cells into the cell cycle by α-factor removal showed low levels of phosphorylated (active) Kss1p early in the cell cycle (**Fig. 1B**, P~Kss1, 0 min to 80 min, G1, S and G2). When the cyclin Clb2p-Myc levels dropped in M phase (Richardson *et al.* 1992; Irniger *et al.* 1995; Wasch and Cross 2002; Cross *et al.* 2005; Eluere *et al.* 2007; Kuczera *et al.* 2010; Cepeda-garcia 2017), P~Kss1p levels increased (**Fig. 1B**, 100 min). Although the cell synchronization deteriorated in the second cycle (based on the failure of Clb2p-Myc to fully disappear), P~Kss1p levels decreased (at 140 min) and then increased again (at 160 min), presumably as cells progressed through the second cell cycle. An increase in P~Kss1p levels in the second cell cycle was also observed (from 17 to 24), which may be due to nutrient depletion at this point in the culture-growth cycle. Activation of the mating pathway might cause a reduction in fMAPK pathway activity through a variety of mechanisms, such as Tec1 degradation (Bao *et al.* 2004; BrÜckner *et al.* 2004; Chou *et al.* 2004) or altered *TEC1* levels by the cell-cycle transcriptional regulator Swi5 (Spellman *et al.* 1998). In an independent approach, cells were synchronized by hydroxyurea (HU), which arrests cells in S phase (Slater 1973; Slater 1974; KoÇ *et al.* 2004). Release of cells from HU treatment also showed low P~Kss1p levels that increased as cells progressed through the cell cycle (*Fig. S1A*) In this case, P~Kss1p levels increased prior to the disappearance of Clb2-Myc levels. These results demonstrate that the activity of the fMAPK pathway fluctuates throughout the cell-cycle.

The fMAPK pathway shares components with the Ste11p-branch of the HOG pathway (**Fig. 1A**). To determine whether the activity of the HOG pathway changes throughout the cell cycle, the phosphorylation of the MAP kinase Hog1p was measured in cells lacking the Sln1p branch (**Fig. 1C**, *ssk1*Δ). As previously reported (Posas and Saito 1997), asynchronous *ssk1*Δ cells exposed to 1M sorbitol showed elevated P~Hog1p levels (**Fig. 1C**, P~Hog1, Async + 1M Sorb). Synchronized *ssk1*Δ cells exposed to 1M sorbitol also showed elevated P~Hog1p levels (**Fig. 1C**, P~Hog1, YEPD + 1M Sorb, 0, 30, 60, and 90 min). In synchronized cells not exposed to osmotic stress, basal P~Hog1p levels showed some periodicity, but compared to P~Kss1p, became reduced throughout the cell cycle (*Fig. S1B).* These results indicate that the HOG pathway can be activated at any point in the cell cycle.

Cells lacking an intact HOG pathway (*pbs2*Δ or *hog1*Δ mutants) exposed to osmotic stress exhibit crosstalk to the mating (O’Rourke and Herskowitz 1998) and fMAPK pathways [**Fig. 1A**, orange arrow (Pitoniak *et al.* 2009)]. In cells lacking Pbs2p, the fMAPK pathway showed elevated P~Kss1p levels in response to 1M sorbitol (**Fig. 1C**, P~Kss1, *pbs2*Δ, YEPD + 1M Sorb). Because P~Kss1p levels were high at all time points tested (0, 30, 60, and 90 min), the cell-cycle regulation of the fMAPK pathway appears to have been lost in the *pbs2*Δ mutant. P~Kss1p levels were also elevated to some degree in *PBS2*+ cells exposed to osmotic stress (**Fig. 1C**, P~Kss1, *ssk1*Δ, YEPD + 1M Sorb at 30, 60, and 90 min but not 0 min), which might result from basal crosstalk between the pathways (Hao *et al.* 2008). As expected, P~Hog1p was not detected in the *pbs2*Δ mutant exposed to sorbitol, because Pbs2p is required to phosphorylate Hog1p. Therefore, during crosstalk, the fMAPK pathway can be activated at early stages in the cell cycle. One implication of this result is that shared components between HOG and fMAPK pathways are present and have the capacity to function at early stages in the cell cycle.

### Cell-Cycle Regulation of the fMAPK Pathway Occurs at the Level of Msb2p and Sho1p: MSB2 Expression is Cell-Cycle Regulated

To define how the fMAPK pathway is regulated throughout the cell cycle, genetic suppression analysis was performed. Genetic suppression analysis can order components in a pathway using gain- and loss-of-function alleles. Hyperactive versions of fMAPK pathway components, Msb2p* [GFP-Msb2p, (Adhikari *et al.* 2015b)], Sho1p^P120L^(Vadaie *et al.* 2008), and Ste11p-4 (Stevenson *et al.* 1992) were examined for the ability to bypass the low levels of fMAPK pathway activity seen early in the cell cycle. Ste11p-4 bypassed the low levels of fMAPK pathway activity (**Fig. 2A**). By comparison, Msb2p* and Sho1p^P120L^ showed a partial bypass (*Fig. S2, A-C).* These results indicate that the cell cycle regulation of the fMAPK pathway occurs above Ste11p and at the level of Msb2p and Sho1p in the fMAPK cascade.

**Figure 2.**
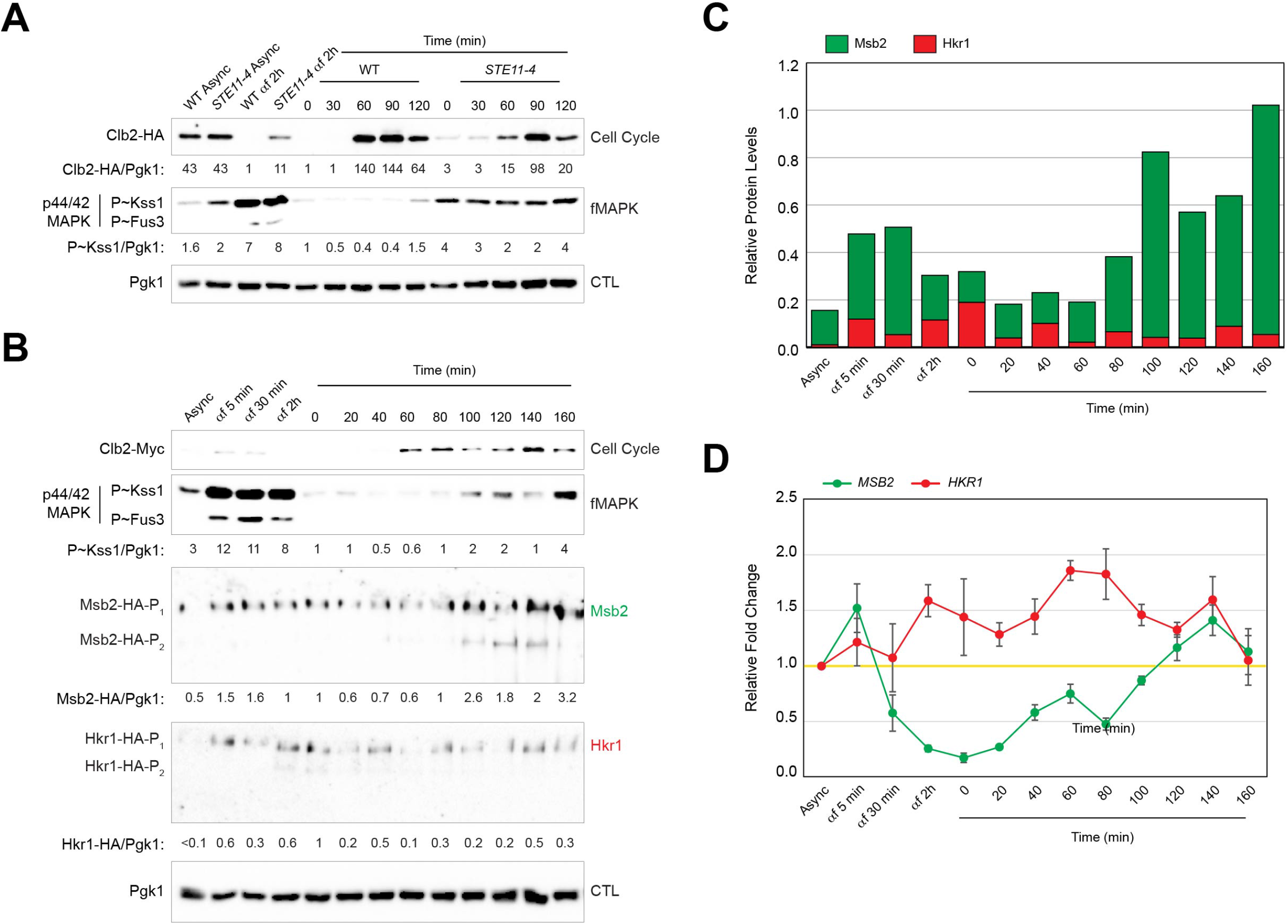
Cell-cycle pattern of fMAPK activity corresponds to different expression patterns of the signaling mucins, Msb2p and Hkr1p. (**A**) Immunoblot analysis of synchronized cultures of wild type (PC7492) and *STE11-4* strain (PC7492 + p*STE11-4′*) released in YEPD. See Figure 1B for details. Numbers refer to the ratio of P~Kss1p to Pgk1p relative to wild type at time 0, which was set to 1. WT, wild type. CTL, loading control. (**B**) Immunoblot analysis of wild-type cells containing Msb2p-HA (PC7495) or Hkr1p-HA (PC7602) synchronized in G1 by α factor and released in YEPD medium. Samples were harvested and probed as in Figure 1B. Numbers refer to the ratio of P~Kss1p to Pgk1p levels, Msb2p-HA to Pgk1p levels, and Hkr1p-HA to Pgk1p levels relative to time 0, which was set to 1. (**C**) Quantitation of Msb2p-HA and Hkr1p-HA protein levels from panel B. Stacked bar graph with protein levels relative to Pgk1p levels are shown. Green bar, Msb2p levels; red bar, Hkr1p levels. (**D**) RT-qPCR analysis of wild-type cells (PC7602) synchronized in G1 by *α*-factor arrest and released in YEPD medium. Fold changes in the mRNA levels of *MSB2* (green) and *HKR1* (red) at indicated time points relative to the asynchronous culture (Async). Yellow line, asynchronous levels. Error bars represent standard error of mean (S.E.M) between 2 biological replicates. One-way ANOVA with Dunnett’s test was used for statistical analysis. The fluctuations in *MSB2* mRNA levels were statistically significant (p-value < 0.01). Specifically, αf = 5min, αf = 2h, t=0, t= 20 min, and t= 80 min were different from the asynchronous and other time points. The fluctuations seen in *HKR1* mRNA levels were not statistically significant (p-value = 0.55, significance < 0.05).

We also noticed that although wild-type cells showed low Clb2p-HA levels after 2 h of α-factor treatment (**Fig. 2A**, Clb2-HA, wT αf 2h) indicative of complete arrest, cells containing Ste11p-4 showed elevated Clb2p-HA levels (**Fig. 2A**, Clb2-HA, *STE11-4* αf 2h), which is suggestive of a delay in G1 arrest. The delay might result from elevated fMAPK activity, which delays the cell cycle in G1 (Loeb *et al.* 1999; Madhani *et al.* 1999) and G2/M (Ahn *et al.* 1999; Rua *et al.* 2001; Vandermeulen and Cullen 2020). In line with this possibility, cells containing Ste11p-4 also showed a delay in accumulation in Clb2p-HA (**Fig. 2A**, compare WT at 60 min with *STE11-4* at 60 min; see also **Fig. 1C**, compare Clb2-HA levels in WT [*ssk1*Δ] and *pbs2*Δ at 60 and 90 min).

The activity of the fMAPK pathway might fluctuate if the levels of one or more components change throughout the cell cycle. We first examined Msb2p levels, which directly impact fMAPK pathway activity (Cullen *et al.* 2004) and drive induction of fMAPK pathway targets (Pitoniak *et al.* 2009). Examining the level of a functional Msb2p-HA protein, expressed under the control of the *MSB2* promoter at its endogenous locus in the genome, showed low levels early in the cell cycle, which peaked later in the cell cycle (**Fig. 2B**, Msb2) Msb2p-HA levels showed a similar pattern in cells synchronized by HU (*Fig. S1A*).

By comparison, the levels of the HOG pathway mucin, Hkr1p, (Tatebayashi *et al.* 2007; Pitoniak *et al.* 2009; Yang *et al.* 2009) also expressed as an HA fusion from its endogenous promoter in the genome, did not show an increase throughout the cell cycle (**Fig. 2B**, Hkr1). Because both mucins contained the same epitope fusion (HA) internal to both proteins in their glycosylated extracellular domains, the levels were able to be compared. Msb2p-HA levels were 15-fold higher than Hkr1p-HA levels in asynchronous (**Fig. 2C**, Async) and synchronized cultures (**Fig. 2C**, 100 and 160 min). Comparative proteomic studies show a similar trend (Breker *et al.* 2013; Yofe *et al.* 2016). The relative abundance of the mucins might impact the activities of the fMAPK and the HOG pathways throughout the cell cycle.

One way that protein levels are regulated is by changes in gene expression. To test whether the levels of Msb2p and Hkr1p result from different patterns of gene expression, mRNA levels of *MSB2* and *HKR1* were examined throughout the cell cycle. *MSB2* mRNA levels were low following release from α-factor treatment and higher at cells progressed through the cell cycle (**Fig. 2D**, *MSB2*). By comparison, *HKR1* mRNA levels were high early in the cell cycle (**Fig. 2D**, *HKR1*) but did not otherwise fluctuate. Thus, *MSB2* and *HKR1* genes show different patterns of gene expression throughout the cell cycle. Moreover, the *MSB2* expression was 5-fold higher than *HKR1* based on analysis of a previously published dataset that examined the levels of *MSB2-lacZ* and *HKR1-lacZ* fusions in asynchronous cells (Pitoniak *et al.* 2009).

### Comparing fMAPK Pathway Activity Under Basal and Pathway-Inducing Conditions Throughout the Cell Cycle

Since Msb2p regulates the fMAPK pathway through positive feedback, Msb2p levels might precede, and therefore induce P~Kss1p levels at M/G1. Alternatively, *MSB2* expression might be induced after P~Kss1p induction, because *MSB2* is a target of the pathway. To determine if the rise in Msb2p levels precede Kss1p activation, the levels of Msb2p-HA and P~Kss1p were compared following release from α factor at short time intervals. Low levels of Msb2p-HA at the beginning of the cell cycle gradually increased and peaked by 40-50 min, which was prior to the increase in P~Kss1p levels (**Fig. 3A**). Although Msb2p levels are known to directly regulate fMAPK activity (ref-Cullen 2004, Hema and Nadia), the fact that increase at 40-50 min did not significantly increase P~Kss1p levels. There may be another regulation at this stage (G1/S boundary) preventing Kss1p activation, such as spatial regulation of Sho1p (see below). After a drop at 80 min, Msb2p-HA levels increased again to peak at 100 min, when P~Kss1p levels rose at M/G1 (**Fig. 3A**, P~Kss1). These fluctuations correlated with *MSB2* mRNA levels (see **Fig. 2D**). Thus, Msb2p protein levels rise prior to the rise in P~Kss1p levels.

**Figure 3.**
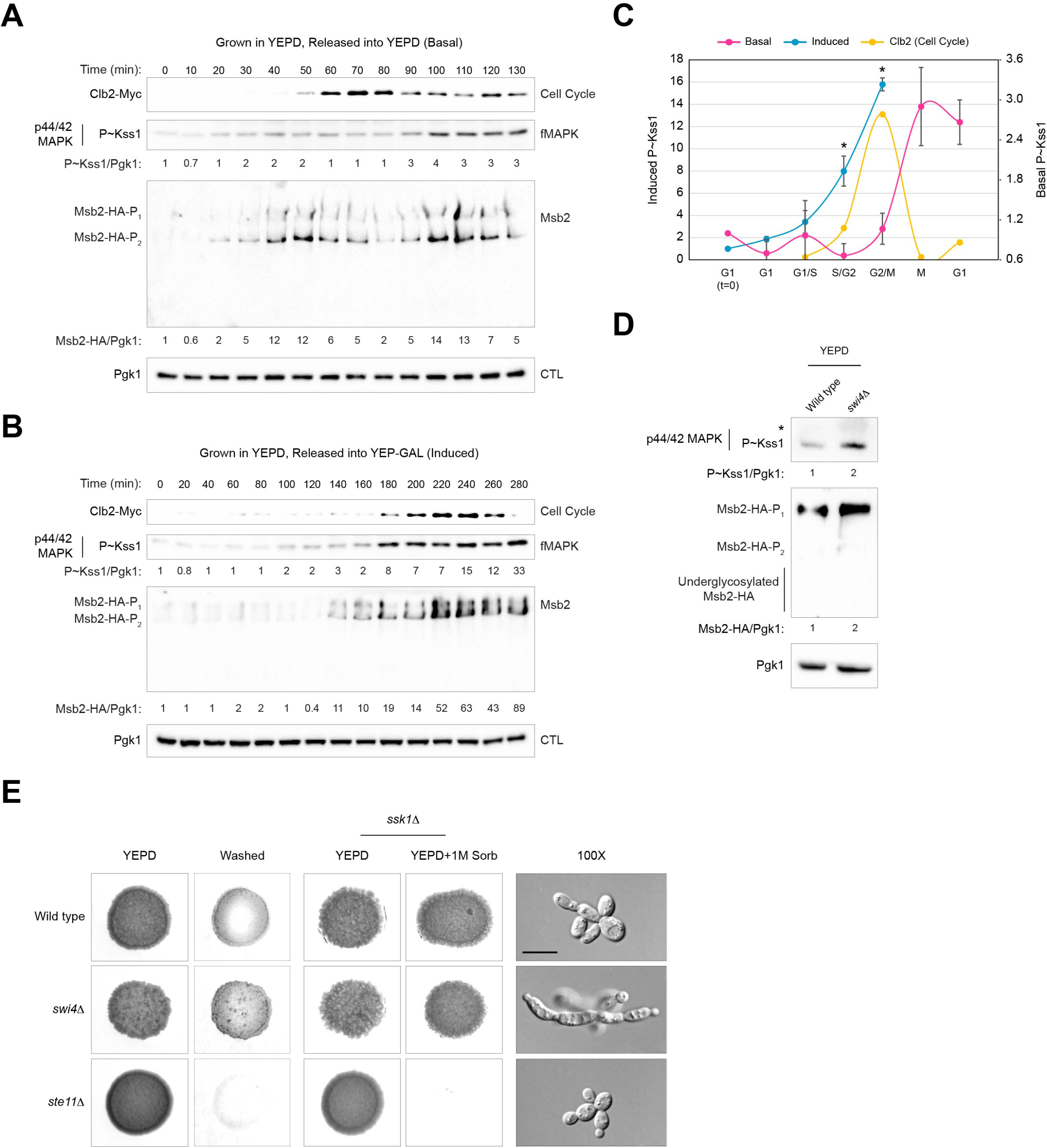
Cell-cycle regulation of the fMAPK pathway is impacted by the activation state of the pathway and the G1/S transcription factor, Swi4p. (**A**) Immunoblot analysis of wild-type cells containing Msb2p-HA (PC7495) pre-grown and synchronized in YEPD and released into YEPD medium. Samples were harvested during the release at indicated time points and probed as in **Fig. 2B**. Numbers refer to the ratio of P~Kss1p to Pgk1p levels and Msb2p-HA to Pgk1p levels relative to time 0, which was set to 1. (**B**) Same as panel A except, cells were pre-grown and synchronized in YEPD and released into YEP-GAL medium. (**C**) Quantitation of relative P~Kss1p levels under basal (pink) and induced (blue) conditions. Error bars represent S.E.M from 2 independent trials. For induced, a biological replicate from (Prabhakar *et al.* 2020) was used for analysis. Yellow curve, Clb2p-levels. X-axis, different phases of cell cycle were determined by time points corresponding to Clb2p-levels after the release. For basal conditions, G1 (t=0), t=0; G1, t=20; G1/S, t=40; S/G2, t=60; G2/M, t=80; M, t=100; G1, t=120. For induced conditions, G1 (t=0), t=0; G1, t=100; G1/S, t=160; S/G2, t=200; G2/M, t=240. One-way ANOVA with Tukey’s test was used for statistical analysis. Asterisk, p-value < 0.05. (**D**) Immunoblot analysis of wild type (PC999) and *swi4*Δ (PC3428) in YEPD. Numbers refer to the ratio of P~Kss1p to Pgk1p levels and Msb2p-HA to Pgk1p levels relative to wild type, which was set to 1. (**E**) Left, PWA analysis of wild type (PC999), *swi4*Δ (PC3428), and *ste11*Δ (PC611). Colonies were grown for 2d. Middle, wild type (PC6810), *swi4*Δ (PC7626) and *ste11*Δ (PC2061) were grown on YEPD and YEPD+1M Sorbitol (Sorb) to evaluate HOG pathway response. Colonies were grown for 4d. Right, DIC images of wild type (PC6810), *swi4*Δ (PC7626) and *ste11*Δ (PC2061) grown in YEPD. Scale, 10 microns.

To this point, fMAPK pathway activity was examined under basal or non-inducing conditions. Nutrient limitation, including nitrogen (Gimeno *et al.* 1992) and carbon (Cullen and Sprague 2000) can trigger filamentous growth, which is thought to be a type of nutrient foraging response seen in fungi. The fMAPK pathway is induced by growth in the non-preferred carbon source galactose [YEP-GAL (Karunanithi and Cullen 2012)]. Cells grown in YEPD and synchronized by α factor that were released into YEP-GAL medium (inducing conditions) showed a delay in P~Kss1p accumulation, which indicates that the fMAPK pathway is cell-cycle regulated under inducing conditions (**Fig. 3B**), consistent with our previous observations (Prabhakar *et al.* 2020). Msb2p-HA levels also accumulated in YEP-GAL prior to the increase in P~Kss1p levels (**Fig. 3B**). Thus, the increase in Msb2p levels precedes and might therefore contribute to the accumulation of P~Kss1p levels throughout the cell cycle.

Clb2p accumulation showed an extended delay in YEP-GAL (**Fig. 3B**, 180 min) compared to YEPD media (**Fig. 3A**, 60-80 min). The extended delay in cell-cycle progression in YEP-GAL might result from glucose repression. Glucose repression involves the transcriptional repression of genes (*GAL* genes and many other genes) that metabolize non-preferred carbon sources (Carlson and Botstein 1982; Nehlin *et al.* 1991; Wilson *et al.* 1996; De Vit *et al.* 1997). To examine fMAPK pathway activity in response to a sustained induction, cells pregrown in YEP-GAL were synchronized and monitored for fMAPK activity. These cells also showed low levels of P~Kss1p after release from α-factor arrest (*Fig. S3*), but not the extended delay seen under pathway-inducing conditions. Thus, cell-cycle regulation of the fMAPK pathway is seen in basal, induced, and sustained-inducing conditions.

In YEP-GAL media, a new pattern of cell-cycle regulation was observed. Compared to basal conditions, where P~Kss1p levels peaked after Clb2p-HA levels fell (**Fig. 3A**), under inducing conditions, P~Kss1p levels peaked at the same time as Clb2p-HA accumulation (**Fig. 3B**). A similar trend was seen under sustained-inducing conditions (*Fig. S3).* Graphing P~Kss1p induction under the two conditions showed the differences in timing between basal (**Fig. 3C**, pink) and inducing (blue) conditions, compared to Clb2p-HA levels (yellow). The ability of the fMAPK pathway to partially bypass the cell-cycle regulation may result from GAL-dependent induction of the fMAPK pathway, which could occur by several mechanism including elevated proteolytic processing of Msb2p, which liberates an inhibitory glycodomain (Vadaie *et al.* 2008). As expected from previous observations, P~Kss1p levels were 12-fold higher under inducing conditions than under basal conditions (**Fig 3C**, compare left and right axes).

### Altering Cell Cycle Regulation Impacts fMAPK Pathway Activity and Filamentous Growth

Cell-cycle progression is regulated by transcription factors. One set of transcription factors, SBF (Swi4/6 cell cycle box Binding Factor), induces transcription of genes required for the progression from G1 to S phase (Andrews and Herskowitz 1989; Nasmyth and Dirick 1991; Sidorova and Breeden 1993). We tested whether altering the normal cell-cycle progression, by loss of Swi4p and Swi6p, impacts fMAPK pathway activity and filamentous growth. The *swi4*Δ mutant had elevated levels of P~Kss1p compared to wild-type cells (**Fig. 3D**, P~Kss1). The *swi6*Δ mutant had a severe growth defect and was not tested further. The *swi4*Δ mutant also had elevated levels of Msb2p-HA (**Fig. 3D**, Msb2-HA), which may account for the elevated levels of fMAPK activity. Evaluation of the *swi4*Δ mutant by the plate washing assay (PWA), which measures invasive growth as a readout of fMAPK pathway activity (Roberts and Fink 1994), showed hyper-invasive growth compared to wild-type cells (**Fig. 3E**, washed). The *swi4*Δ mutant was not required for growth on high-osmolarity media (**Fig. 3E**, YEPD+1M Sorb). The *swi4*Δ mutant cells had an elongated appearance (**Fig. 3E**, 100X), which may result from hyperpolarized growth due to an extension of the G1 phase of the cell cycle (White *et al.* 2009) and/or increased fMAPK activity (**Fig. 3D**). Although the *swi4*Δ mutant responded to α factor, the cells did not synchronize, which prevented evaluation of P~Kss1p levels throughout the cell cycle in this mutant. These results indicate that normal cell-cycle progression is required for proper fMAPK pathway activity and filamentous growth.

### Positive Feedback of the fMAPK Pathway Induces the HOG Pathway

The cell-cycle regulation of the fMAPK pathway might extend to transcriptional targets of the pathway. The fMAPK pathway regulates multiple target genes that encode proteins that bring about filamentous growth (Madhani *et al.* 1999; Roberts *et al.* 2000; Vandermeulen and Cullen 2020). A major target of the fMAPK pathway is the gene encoding the cell adhesion molecule Flo11p (Rupp *et al.* 1999; Vinod *et al.* 2008), which promotes cell adhesion during filamentous/invasive growth (Rupp *et al.* 1999). By qPCR analysis, *FLO11* expression showed a similar periodicity as *MSB2*, which indicates that its expression is regulated throughout the cell cycle (**Fig. 4A**). Several transcriptional targets of the fMAPK pathway encode components of the pathway and are induced by positive feedback. The gene encoding the MAP kinase Kss1p (Roberts *et al.* 2000) (**Fig. 4B**) and the TEA/ATS type transcription factor Tec1p (KÖhler *et al.* 2002) (**Fig. 4C**) showed a similar pattern of cell-cycle regulation as *MSB2.* By comparison, the transcription factor Ste12p, which functions in the mating and the fMAPK pathways (Roberts and Fink 1994) showed a different pattern of expression, perhaps because its expression is induced by pheromone (**Fig. 4D**). Therefore, many of the transcriptional targets of the fMAPK pathway that we tested showed cell-cycle regulated expression profiles.

**Figure 4.**
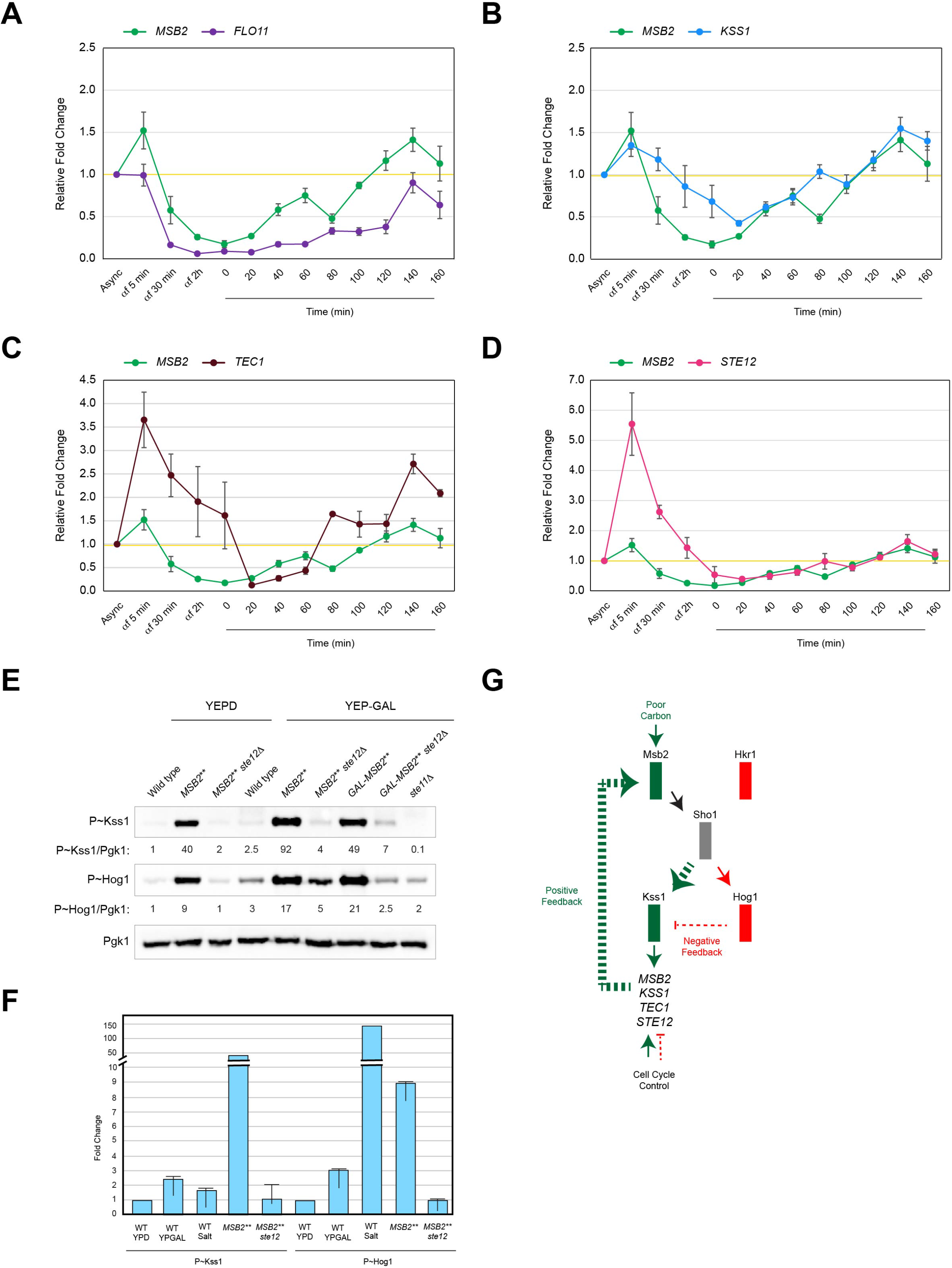
Expression profile of fMAPK components throughout the cell cycle and role of cross-pathway feedback in regulating fMAPK and HOG activities. (**A**) Fold changes in the mRNA levels of *MSB2* (green) and *FLO11* (purple) at indicated time points in synchronized cells relative to the asynchronous culture (Async). See **Fig. 2D** for details. *MSB2* values were taken from **Fig. 2D**. One-way ANOVA with Dunnett’s test was used for statistical analysis. The fluctuations in *FLO11* mRNA levels were statistically significant (p-value < 0.01). Specifically, *α*f = 30 min, *α*f = 2h, t=0 min-120 min were different from the asynchronous and other time points. (**B**) Same as panel A, except *MSB2* (green) and *KSS1* (blue). The fluctuations in *KSS1* mRNA levels were statistically significant (p-value < 0.01). t= 20 min and t= 140 min were different from the asynchronous and other time points. (**C**) Same as panel A, except *MSB2*(green) and *TEC1* (brown). Based on ANOVA with Dunnett’s statistical test, the fluctuations in *TEC1* mRNA levels at αf = 5 min and at t= 140 min were different from the asynchronous and other time points. (**D**) Same as panel A, except *MSB2* (green) and *STE12* (red). Based on ANOVA with Dunnett’s statistical test, the fluctuations in *STE12* mRNA levels at αf = 5 min and at αf = 30 min were different from the asynchronous and other time points. (**E**) Immunoblot analysis of wild type (PC538), *MSB2\*\** (PC1516), and *MSB2** ste12*Δ (PC1811) grown in YEPD and YEP-GAL and *GAL-MSB2\*\** (PC1806), *GAL-MSB2** ste12*Δ (PC1837), and *ste11*Δ(PC611) grown in YEPG-GAL. Numbers refers to the ratio of P~Kss1p to Pgk1p levels and P~Hog1p to Pgk1p levels relative to wild type in YEPD, which was set to 1. (**F**) Comparison of P~Kss1p levels and P~Hog1p levels in the indicated strains under basal and pathway-inducing conditions for the fMAPK and HOG pathways. The histogram represents the ratio of P~Kss1p to Pgk1p levels and P~Hog1p to Pgk1p levels relative to wild-type, which was set to 1. Error bars represent s.e.m. from 2 independent trials. (**G**) Model showing role of poor carbon source, cell cycle, positive feedback of pathway components and the HOG pathway in regulating the activity of fMAPK pathway. Arrows and proteins shown in green refer to induction; lines and proteins shown in red refer to inhibition; cell cycle has inducing and inhibiting effect depending on the stage of the cell cycle; Sho1p, shared protein. Dashed green arrow refers to preferential stimulation of fMAPK pathway through positive feedback loop that involves Msb2p.

To further explore the relationship between positive feedback and fMAPK pathway activity, we examined the consequences of hyperactivating the fMAPK pathway in cells lacking the transcription factors that control positive feedback. Cells containing another hyperactive version of Msb2p, Msb2p^Δ100-818^, Msb2p**\*\***, hyperactivated the fMAPK pathway through positive feedback, because the transcription factor Ste12p was required [**Fig. 4E**, P~Kss1, (Cullen *et al.* 2004; Vadaie *et al.* 2008; Prabhakar *et al.* 2020)]. Positive feedback occurred in basal (YEPD) and inducing (YEP-GAL) conditions (**Fig. 4E**). Msb2p** expressed from an fMAPK-independent promoter also required Ste12p to hyperactive the fMAPK pathway [**Fig. 4E**, P~Kss1, GAL-*MSB2** ste12*Δ,(Prabhakar *et al.* 2020)], which indicates that in addition to *MSB2* other components of the pathway (like Kss1p and Tec1p) are required for positive feedback.

We also examined the activity of the HOG pathway under this condition. In addition to osmotic stress, the HOG pathway can also be induced by the non-preferred carbon source, galactose (Adhikari and Cullen 2014). Unexpectedly, we found that Msb2p** also stimulated the HOG pathway in a Ste12p-dependent manner (**Fig. 4E**, P~Hog1). The effect was seen in basal and pathway-inducing conditions (**Fig. 4E**). This result indicates that positive feedback from the fMAPK pathway leads to activation of the HOG activity. Given that the HOG pathway functions antagonistically to the fMAPK pathway (Davenport *et al.* 1999; Adhikari and Cullen 2014), these results might describe a novel type of cross-pathway feedback, where positive feedback through one pathway (fMAPK) induces another pathway to modulate its activity. It might be interesting to note that positive feedback through the fMAPK pathway was higher (20-fold) than positive feedback through the HOG pathway (9-fold in GLU and 3-fold in GAL, **Fig. 4F**). Other measurements also showed key similarities and differences between the pathways. Thus, Msb2p induces both MAP kinase pathways at different levels to produce an appropriate response. This feedback regulation can be described by a simple model, which shows the relationship between the two pathways (**Fig. 4G**). In this relationship, Msb2p induces one pathway at high levels (fMAPK, green), and its antagonistic pathway to lower levels (HOG, red), resulting in a modulated response.

### Sho1p Levels and Localization Change Throughout the Cell Cycle

To regulate the fMAPK pathway, Msb2p interacts with the tetraspan protein, Sho1p (O’Rourke and Herskowitz 1998; Cullen *et al.* 2004). The levels and expression of Sho1p were also examined. Sho1p-GFP levels rose prior to accumulation in Clb2p-Myc levels in G2/M (**Fig. 5A**) and dropped when P~Kss1p levels increased. The drop in Sho1p-GFP levels corresponding to fMAPK pathway activation might be due to turnover of active Sho1p. Specifically, a hyperactive version of Sho1p, Sho1p^P120L^, shows elevated turnover compared to the wild-type protein (Adhikari *et al.* 2015a). *SHO1* mRNA showed a similar pattern of cell-cycle regulation (**Fig. 5B**, orange line). The increase in *SHO1* mRNA levels correlated with the increase in protein levels. Incidentally, Sho1p-GFP protein levels increased after 30 min of α-factor treatment (**Fig. 5A**, Sho1-GFP), which might occur because Sho1p has a function in mating (Nelson *et al.* 2004). Therefore, genes encoding two of the sensors for the fMAPK pathway, *MSB2* and *SHO1*,show similar patterns of cell-cycle regulated gene expression.

**Figure 5.**
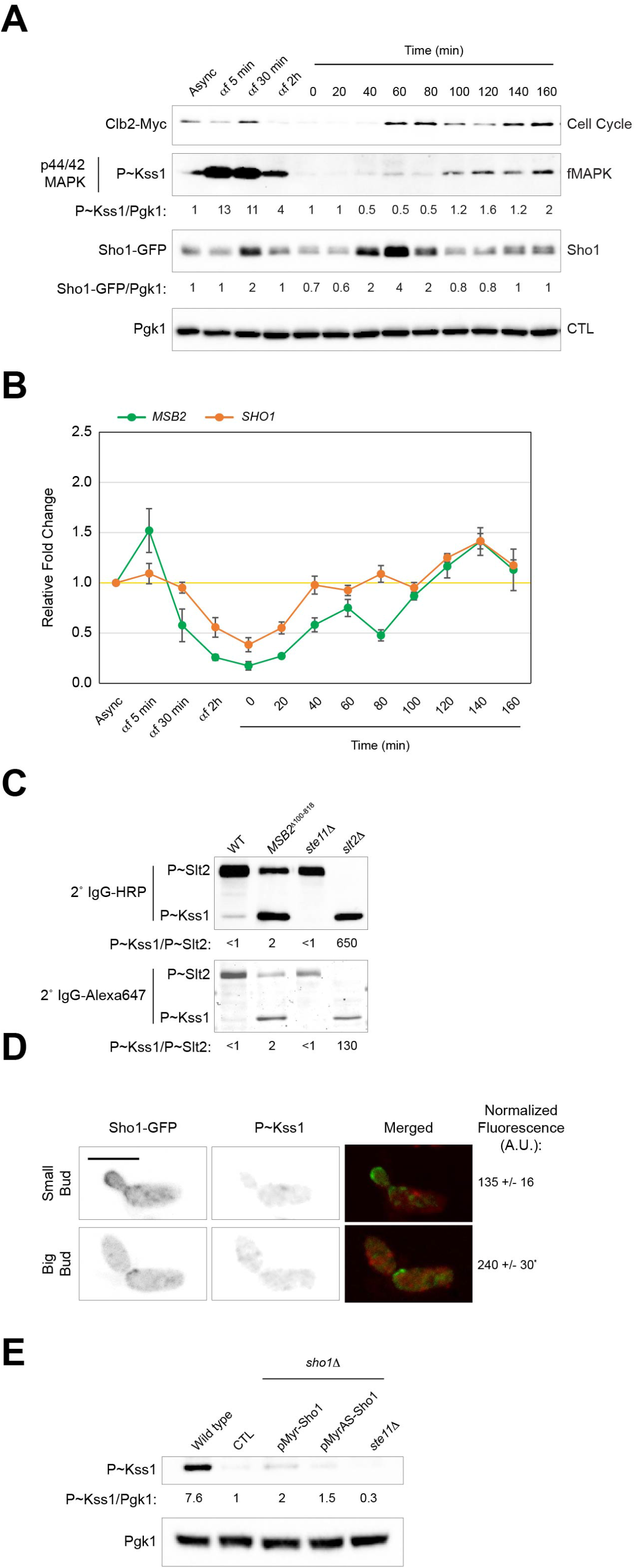
Expression and localization of Sho1p throughout the cell cycle. (**A**) Immunoblot analysis of wild-type cells containing Sho1p-GFP (PC7495 + p*SHO1-GFP::NAT*) synchronized in G1 by α-factor arrest and released in YEPD. Samples were harvested and probed as in Figure 1B. Numbers refer to the ratio of P~Kss1p to Pgk1p levels and Sho1p-GFP to Pgk1p levels relative to time 0, which was set to 1. (**B**) Fold change in the mRNA levels of *MSB2* (green) and *SHO1* (orange) at indicated time points in synchronized cells relative to the asynchronous culture (Async). See **Fig. 2D** for details. *MSB2* values were taken from **Fig.2D**. Based on ANOVA with Dunnett’s statistical test, the fluctuations in *SHO1* mRNA levels at αf = 2h, t= 0 min, t= 20 min, and t=140 min were different from the asynchronous and other time points. (**C**) Immunoblot analysis of wild type (PC538), *MSB2\*\** (PC1516), *ste11*Δ (PC611) and *slt2*Δ (PC3394) using anti-p44/42 rabbit primary antibody and anti-rabbit goat HRP conjugated or anti-rabbit goat Alexa 647 as secondary antibody. Numbers refer to ratio of P~Kss1p to P~Slt2p levels. (**D**) indirect immunofluorescence of *slt2*Δ cells containing Sho1p-GFP (PC3394 + p*SHO1-GFP::NAT).* Representative cells where Sho1p-GFP is present in the bud cortex (small bud) or at the mother-bud neck (large bud) are shown. Mid-log cells grown in YEP-GAL were stained with anti-p44/42 rabbit primary antibody followed by anti-rabbit goat Alexa 647 secondary antibody. Numbers refer to normalized pixel intensity of P~Kss1p in each cell. Error represents S.E.M among 15 cells. Two-sided *t* test was used for statistical analysis. Asterisk, intensity values in large buds were different (p-value < 0.05) from those in small buds. A. U., arbitrary units. Scale bar, 5 microns. (**E**) Immunoblot analysis of wild-type cells (PC538), *sho1*Δ (PC1531), *sho1*Δ containing pMyr-Sho1 and pMyrAS-Sho1 plasmids, and *ste11*Δ (PC611). Numbers refer to the ratio of P~Kss1p to Pgk1p levels relative to *sho1*Δ (CTL), which was set to 1. CTL, control.

Although the Msb2p, Sho1p, and Opy2p proteins form a complex in the plasma membrane (Tatebayashi *et al.* 2015; Yamamoto *et al.* 2016), they have different patterns of localization and turnover (Adhikari *et al.* 2015a). Processed Msb2p is turned over by the E3 ubiquitin ligase Rsp5p and is mainly localized to the lysosome/vacuole (Adhikari *et al.* 2015a; Adhikari *et al.* 2015b; Prabhakar *et al.* 2019), whereas Sho1p primarily localizes to plasma membrane (Raitt *et al.* 2000; Reiser *et al.* 2003; Pitoniak *et al.* 2009) at the growth tip in developing buds and the mother-bud neck in large buds. To better understand the contribution of Msb2p and Sho1p in the cell-cycle regulation of the fMAPK pathway, the localization of the Msb2p-GFP and Sho1p-GFP proteins were examined as cells progressed through the cell cycle. Time-lapse fluorescence microscopy was performed by co-localization of GFP fusion proteins to Msb2p and Sho1p, and the septin and the cell-cycle marker, Cdc3p-mCherry (Kim *et al.* 1991; Lippincott *et al.* 2001). Due to its high turnover rate (Adhikari *et al.* 2015a; Adhikari *et al.* 2015b), Msb2p-GFP showed a predominately vacuolar localization pattern. Although Msb2p-GFP was at the periphery in some cells, its localization was not otherwise informative (<u>Movie 1</u>). By comparison, Sho1p-GFP localized to different parts of the cell throughout the cell cycle, including presumptive bud sites, the tip of developing buds and the mother-bud neck (<u>Movies 2 and 3</u>). The same pattern was seen under inducing conditions (Gal), except that Sho1p-GFP was polarized at the distal pole for an extended period (<u>Movie 4</u>). In cells grown in Gal, which bud distally, it was clear that Sho1p-GFP was localized to the at the mother-bud neck during prior to cytokinesis, when the septin ring splits into a double ring [<u>Movie 4</u> (Kim *et al.* 1991; Lippincott *et al.* 2001; Bi and Park 2012)].

Because Sho1p was localized to the mother-bud neck during septin ring split, it appeared that Sho1p-GFP was at the mother-bud neck during the period of the cell cycle when cells experienced elevated fMAPK pathway activity. To further define the location of Sho1p during the induction of fMAPK pathway activity, the same cells harvested for immunoblot analysis (in **Fig. 5A**) were examined by microscopy for Sho1p-GFP localization. In synchronized cells, the increase in P~Kss1p levels (**Fig. 5A**, P~Kss1, 100 min) corresponded to an increase in the percentage of cells that showed Sho1p-GFP localization at the mother-bud neck. Synchronized cells containing Sho1p-GFP and Cdc3p-mCherry showed co-localization at the mother-bud neck during the time of elevated fMAPK pathway activity (*Fig. S4A*, 100 min, 45%).

A challenge connecting Sho1p localization to fMAPK pathway activity is that protein localization is evaluated by microscopy, whereas MAPK activity is evaluated by phospho-immunoblot analysis. In mammalian cells, the localization of P~ERK has been evaluated by immunofluorescence (IF), which has revealed insights into the spatial and temporal nature of MAPK pathway signaling (Shapiro *et al.* 1998; Ingram *et al.* 2000; Molgaard *et al.* 2016). Antibodies that detect phosphorylated mammalian ERK also detect the phosphorylated forms of three yeast ERK-type MAPK kinases: Slt2p, for cell wall integrity pathway (Lee *et al.* 1993)], Kss1p, for fMAPK (Cook *et al.* 1997)]; and Fus3p, for the mating pathway (Elion *et al.* 1993)]. Therefore, a problem with iF of P~Kss1p is interference by other P~ERK type MAPKs. To circumvent this problem, the *SLT2* gene was disrupted. In the *slt2*Δ mutant, P~Kss1p was the main band detected using a phospho-MAPK specific antibody that preferentially detects P~Kss1p over P~Fus3p (**Fig. 5C**, *slt2*Δ) As expected, by immunoblot analysis P~Kss1p levels were higher in cells carrying the hyperactive *MSB2*^Δ100-818^ mutant and reduced in cells lacking the MAPKKK Ste11p (**Fig. 5C**, *ste11*Δ). An Alexa 647 fluorophore-conjugated secondary antibody (Thermo fisher, Waltham, MA) showed the same pattern by immunoblot (**Fig. 5C**) and detected P~Kss1p by immunofluorescence (*Fig. S4B*). Cells grown in basal conditions showed brighter P~Kss1p levels than the no antibody control (*Fig. S4B*). Cells grown under pathway-inducing conditions (YEP-GAL) showed brighter P~Kss1p than cells grown in basal conditions (*Fig. S4B).* Therefore, IF of P~Kss1p is a feasible method to evaluate P~Kss1p activity.

Like many MAP kinases, mammalian ERK enters the nucleus upon activation (Chen *et al.* 1992; Lenormand *et al.* 1993; PouyssÉgur *et al.* 2002; Zehorai *et al.* 2010). By comparison, Kss1p has a unique regulatory mechanism. Unphosphorylated Kss1p is present in the nucleus in an inhibitory complex with Ste12p, Tec1p, and Dig1p (Bardwell *et al.* 1996; Cook *et al.* 1997; Bardwell *et al.* 1998). Upon phosphorylation by Ste7p, active Kss1p phosphorylates Ste12p, Tec1p and Dig1p and exits the nucleus (Ma *et al.* 1995; Bardwell *et al.* 1998; Pelet 2017). Consistent with this mechanism, P~Kss1p showed a punctate pattern in the cytoplasm (*Fig. S4B*). P~Kss1p levels were next evaluated throughout the cell cycle. P~Kss1p level were higher in M/G1, based on the signal intensity of mitotic and post-mitotic cells where the nucleus was visible in the mother and the daughter cell (*Fig. S4C*, normalized fluorescence). In addition, cells where Sho1p-GFP was localized to the mother-bud neck showed ~2-fold higher P~Kss1p levels than cells where Sho1p-GFP was localized in buds (**Fig. 5D**). These results demonstrate that Sho1p is localized to the mother-bud neck when cells experience elevated fMAPK pathway activity during M/G1, which defines spatial and temporal aspects to the regulation of fMAPK pathway signaling.

To further test whether Sho1p’s localization is critical for its activity in the fMAPK pathway, a version of Sho1p was examined where its cytosolic signaling domain was anchored to the plasma membrane by a myristylation tag [pMyr-Sho1p (Raitt *et al.* 2000)]. pMyr-Sho1 was not able to induce the fMAPK pathway (**Fig. 5E**), although it has been reported to function in HOG (Raitt *et al.* 2000). Similarly, a version that contains a point mutation in the myristylation site (pMyr-ASSho1) was also defective for fMAPK pathway signaling. Thus, based on this preliminary experiment, it appears that Sho1p’s cytosolic domain is not merely a passive scaffold in regulating the fMAPK pathway.

### Cytokinesis Regulatory Protein Hof1p Regulates the fMAPK Pathway

The fact that Sho1p localizes to the mother-bud neck when cells experience elevated fMAPK pathway activity suggests that the mother-bud neck is a relevant site for fMAPK pathway signaling. To further test this possibility, the role of septins was examined. Septins are heterooligomers that form a cytoskeletal ring at the mother-bud neck and control bud emergence, cytokinesis, and mother-daughter asymmetry (Hall 2008; McMurray and Thorner 2009; Bi and Park 2012). Although septins are essential for viability, and temperature-sensitive alleles of septin genes allow evaluation of septin function. The *cdc12-6* mutant shows normal growth at 25°C and is inviable at 37°C. At 30°C, the *cdc12-6* mutant exhibits cytokinesis defects. At 30°C, the *cdc12-6* mutant had a defect in fMAPK activity, based on immunoblot analysis of P~Kss1p levels (**Fig. 6A**). The *cdc12-6* mutant also showed a defect in the activity of the *FUS1-lacZ* reporter (**Fig. 6B**) and the *FUS1-HIS3* reporter (*Fig. S5A*), which in strains lacking an intact mating pathway (*ste4*Δ) shows dependency on the fMAPK pathway (Cullen *et al.* 2004). Sho1p-GFP was also mis-localized in the *cdc12-6* mutant at 30°C (**Fig. 6C**, arrows), which may account for the signaling defect seen in this mutant. The *cdc12-6* mutant also showed a defect in fMAPK pathway activity at 25°C (**Fig. 6B**, *Fig. S5A*), which we have previously reported is due to a bud-site-selection defect (Basu *et al.* 2016). Therefore, proper septin function is required for fMAPK pathway activity and Sho1p localization.

**Figure 6.**
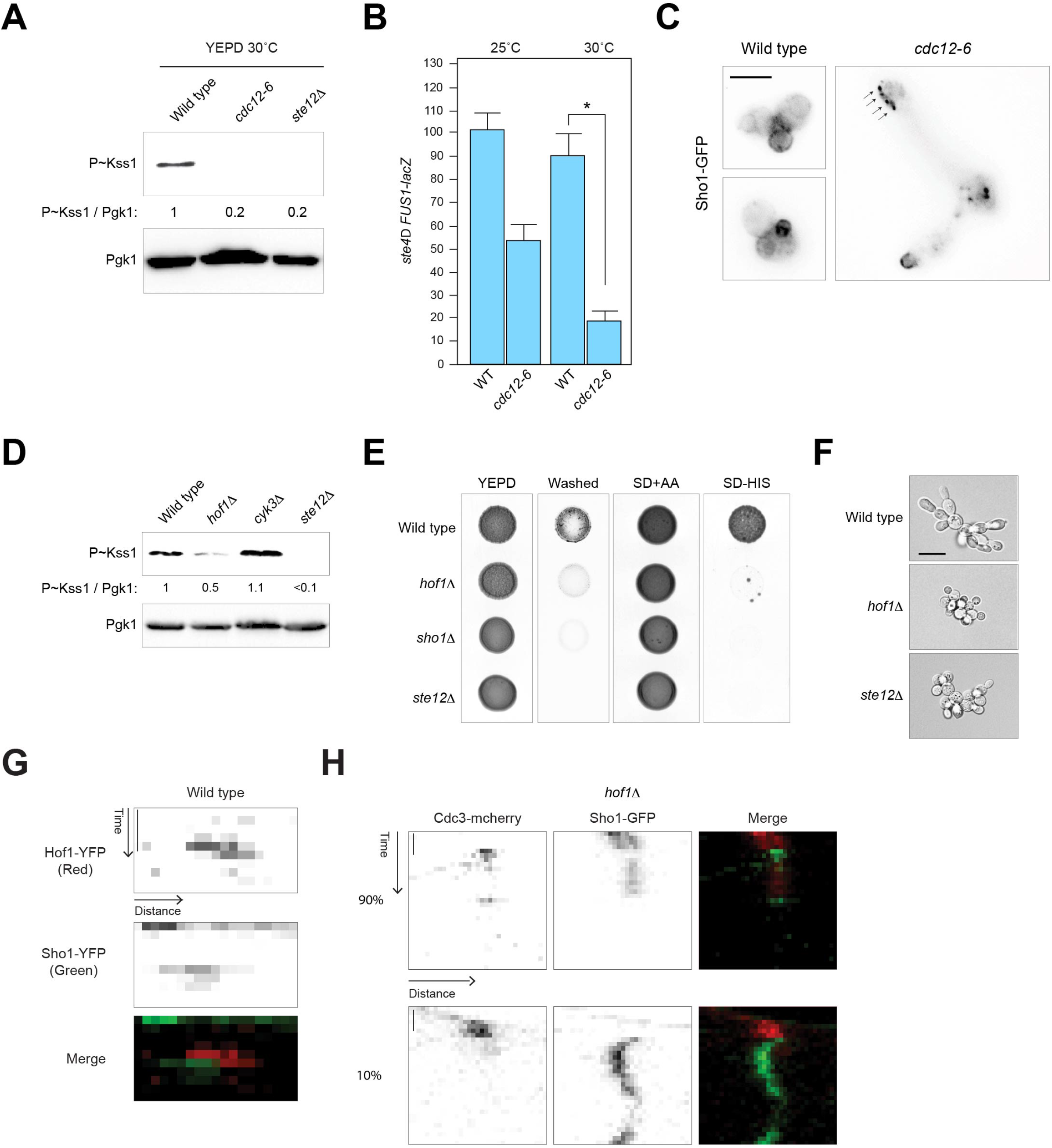
Role of septins and cytokinesis-regulatory proteins in regulating the fMAPK pathway. (**A**) Immunoblot analysis of wild-type cells (PC538), the *cdc12-6* mutant (PC2710), and the *ste12*Δ mutant (PC539) grown in YEPD medium. Numbers refers to the ratio of P~Kss1p to Pgk1p levels relative to wild type, which was set to 1. (**B**) Evaluation of wild-type cells and the *cdc12-6* mutant for fMAPK activity using *FUS1-lacZ* β-galactosidase reporter. Cells were grown to mid-log state at indicated temperatures. (**C**) Wild-type cells and the *cdc12-6* mutant carrying pSho1p-GFP plasmid were grown to mid-log stage in YEPD medium and visualized under GFP channel. (**D**) Immunoblot analysis of wild type (PC538), *hof1*Δ (PC2371), *cyk3*Δ (PC6472), and *ste12*Δ (PC539) strains grown in YEPD medium. See panel A for details. (**E**) The *hof1*Δ mutant was evaluated for invasive growth using plate washing assay (washed) and fMAPK activity using *FUS1*-*HIS3* growth reporter alongside controls. (**F**) Single-cell assay for wild-type cells, the *hof1*Δ mutant and the *ste12*Δ mutant. Diluted cultures of overnight cells were spread onto synthetic medium lacking glucose and incubated at 30°C for 12 h before visualization. Scale bar, 10 microns. (**G**) Hof1p-CFP and Sho1p-YFP localization at the mother-bud neck in wild-type cells (PC2377) represented by a kymograph. Black and white images were false colored red (for Hof1p-CFP) and green (for Sho1p-YFP) in the merged channel to improve visualization. Scale bar, 50 min. (**H**) Kymograph of Sho1p-GFP and Cdc3p-mCherry in the *hof1*Δ mutant (PC7563 + p*SHO1*-*GFP*). Representative kymographs of two types of localization patterns cells are shown. Most cells (90%) show a normal localization pattern (top panels). Some cells (10%) show mis-localization of Sho1p-GFP. Scale bar, 50 min.

At the mother-bud neck, Sho1p interacts with proteins that regulate cytokinesis, including Hof1p, Cyk3p, and Inn1p (Labedzka *et al.* 2012). Hof1p is localized to the mother-bud neck at the initial stages of cytokinesis and moves to the actomyosin ring during cytokinesis (Vallen *et al.* 2000; Meitinger *et al.* 2011; Oh *et al.* 2013). The *HOF1* and *CYK3* genes, and several other genes that regulate aspects of cytokinesis, including *BNI5* and *SHS1*, were disrupted in wild-type strains of the filamentous (Σ1278b) background. The *hof1*Δ mutant, but not the *cyk3Δ, bni5*Δ, or *shsl.\* mutants, showed a defect in fMAPK activity based on P~Kss1 levels and *FUS1-HIS3* activity (**Fig. 6D**and **E**, *Fig. S5B).* Cyk3p might not regulate the fMAPK pathway because that protein does not interact with septins and has a distinct function from Hof1p (Oh *et al.* 2013). The *hof1*Δ mutant was also defective for invasive growth by the plate-washing assay (**Fig. 6E**) and the formation of filamentous cells (**Fig. 6F**) by the single-cell invasive growth assay (Cullen and Sprague 2000). These results establish Hof1p as a regulator of the fMAPK pathway.

Hof1p might regulate the fMAPK pathway by influencing Sho1p localization. Sho1p-YFP and Hof1p-CFP both localized at the mother-bud neck (**Fig. 6G**, kymograph, <u>Movie 5</u>), and Sho1p was mis-localized in the *hof1*Δ mutant (**Fig. 6H**, kymographs; <u>Movie 6</u>, normal cell; <u>Movie 7</u>, abnormal cell), although this phenotype was seen in only ~ 10% of cells, which also showed a cytokinesis defect. Sho1p showed genetic interactions with *HOF1* in that overexpression of *SHO1*, which induces hyperpolarized growth (Vadaie *et al.* 2008; Pitoniak *et al.* 2015), exacerbated the growth defect of the *hof1*Δ mutant (*Fig. S5C).* Hof1p might alternatively impact bud-site-selection. Bud-site-selection proteins that control axial budding localize to the mother-bud neck (Chant *et al.* 1995; Sanders and Herskowitz 1996). Budsite-selection proteins also regulate the fMAPK pathway (Basu *et al.* 2016). The *hof1*Δ mutant had a defect in bud-site selection (*Table 2*, *Fig. S5D*), although the defect was less severe than seen in mutants lacking bud-site-selection proteins (*Table 2*, *bud3*Δ). The budding pattern defect of the *hof1*Δ mutant may not account for its signaling defect, because other cytokinesis regulators that had similar or more severe bud-site-selection defects did not impact fMAPK pathway signaling (Table 2, *Fig. S5D).* Thus, Hof1p may regulate the fMAPK pathway, in part, by regulating the localization of Sho1p.

**Table 1.**
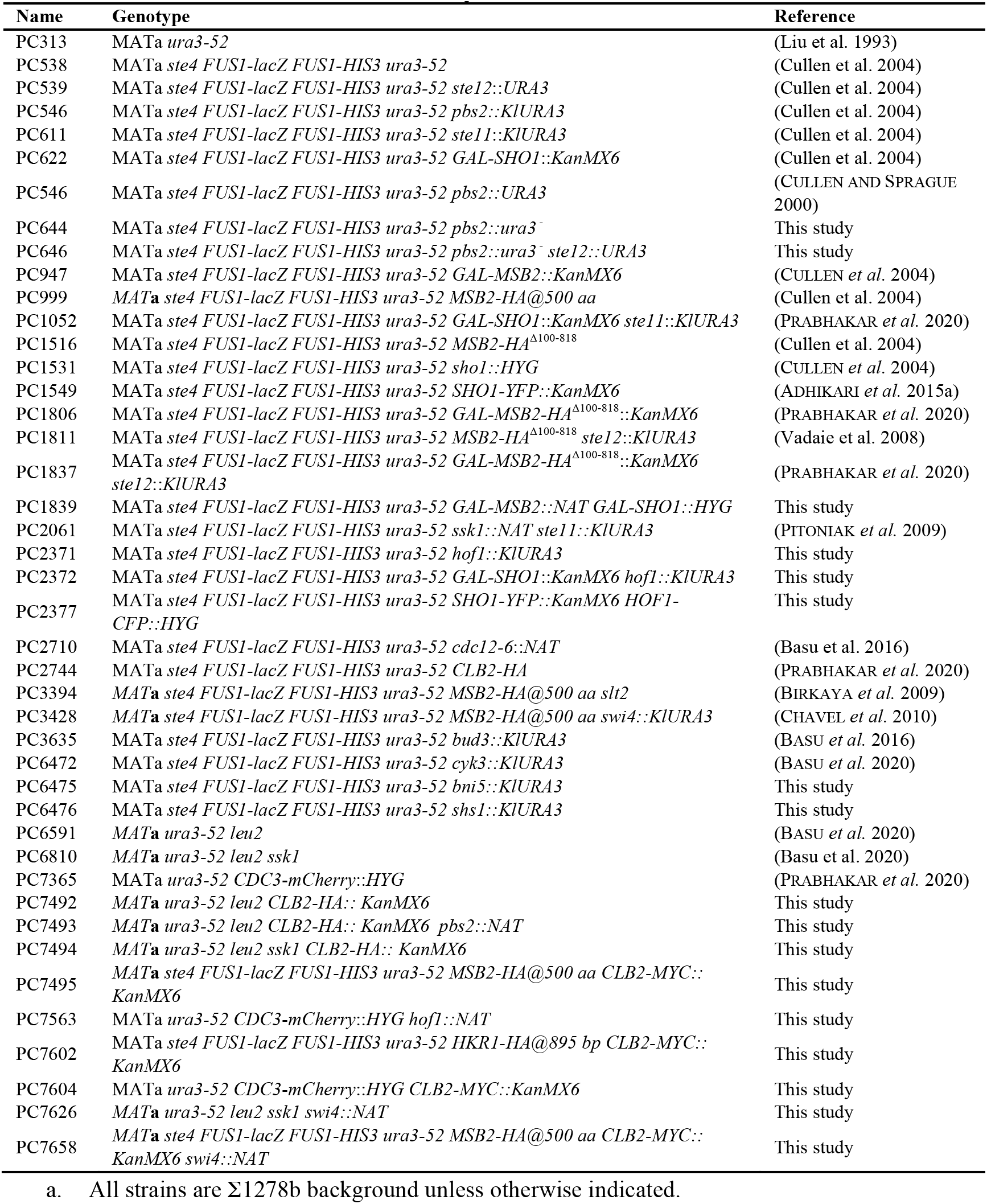
Yeast strains used in the study.

**Table 2.**
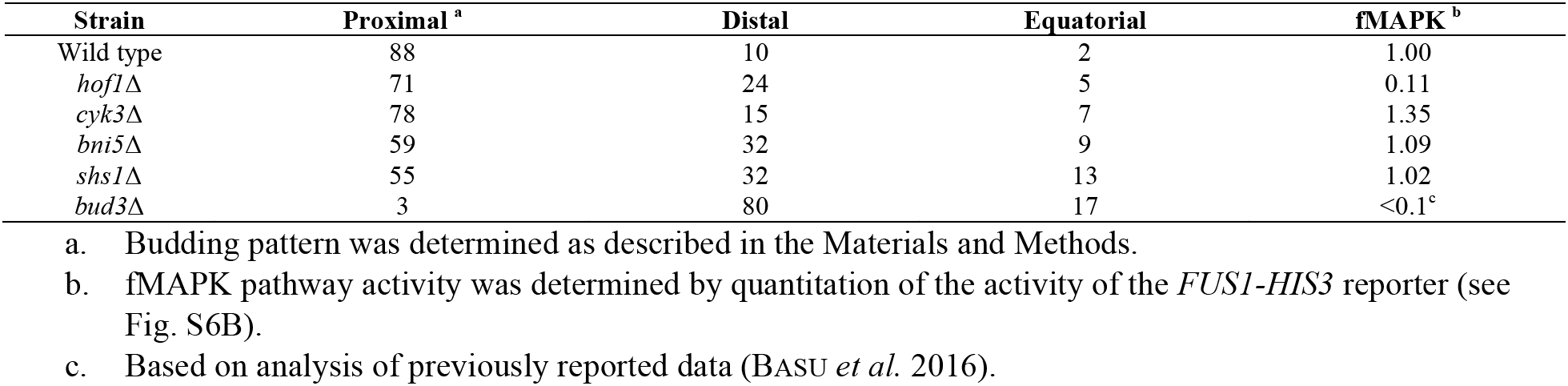
Patterns of bud-site selection for mutants lacking proteins that regulate cytokinesis.

| Strain | Proximal <sup>a</sup> | Distal | Equatorial | fMAPK <sup>b</sup> |
| --- | --- | --- | --- | --- |
| Wild type | 88 | 10 | 2 | 1.00 |
| *hof1*Δ | 71 | 24 | 5 | 0.11 |
| *cyk3*Δ | 78 | 15 | 7 | 1.35 |
| *bni5*Δ | 59 | 32 | 9 | 1.09 |
| *shs1*Δ | 55 | 32 | 13 | 1.02 |
| *bud3*Δ | 3 | 80 | 17 | <0.1<sup>c</sup> |

## DISCUSSION

Signaling pathways are regulated by extrinsic cues, by other pathways in integrated networks, and by spatiotemporal mechanisms to produce a precise signal that generates a physiologically-relevant response. Here, we provide evidence that the MAPK pathway that regulates filamentous growth in yeast is subject to temporal regulation throughout the cell cycle, and spatial regulation by proteins that function at the mother bud neck. We also show that amplification of the fMAPK pathway by positive feedback generates crosstalk to a pathway that shares components, which functions to modulate fMAPK pathway activity. Collectively, these regulatory processes may function to precisely match MAPK pathway activity during a cell differentiation response, as well as provide a mechanism for specific activation of a MAPK pathway that shares components with other MAPK pathways in the same cell type.

### Cell-Cycle Regulation of the fMAPK Pathway

A common function for MAPK pathways is to alter cell-cycle progression. In yeast, MAPK pathways alter the progression of the cell cycle during mating (Strickfaden *et al.* 2007), during filamentous growth (Madhani *et al.* 1999), and in response to osmotic stress (Waltermann *et al.* 2010; Radmaneshfar *et al.* 2013). Mammalian ERKs induce entry into the cell cycle from a G_0_ state (Seger *et al.* 1994) and regulate G1/S (Aktas *et al.* 1997; Leone *et al.* 1997) and G2/M (Wright *et al.* 1999; Dangi *et al.* 2006) transitions including to regulate mitosis (Shapiro *et al.* 1998). By comparison, it is relatively less well understood how the activity of MAPK pathways might themselves be cell-cycle regulated. One reason for this is that cell synchronization experiments, followed by careful measurement of MAPK pathway activity by phospho-immunoblot analysis, or by examining protein localization over the cell cycle by time-lapse fluorescence microscopy are technically challenging especially in complex metazoan systems.

Here we show that the activity of the fMAPK pathway is cell-cycle regulated (*Fig. S6*). The activity of the fMAPK pathway is low in G1, S, and G2, and up in M/G1. By comparison, the HOG pathway can be activated by osmotic stress at any point in the cell cycle. The fact that the HOG pathway is not cell-cycle regulated may not be surprising as cells might be expected to encounter an osmotic stress at any point in their life cycle. One point of regulation occurs at the level of expression of the gene encoding the mucin Msb2p. We also show that the G1/S transcription factor Swi4p regulates the fMAPK pathway activity and filamentous growth. Although Swi4p may regulate the fMAPK pathway in a number of ways, one possibility is by regulating *MSB2* expression. The *MSB2* promoter contains the regulatory element CACGAAA (Breeden and Nasmyth 1987) 466 bp upstream of the start site, which binds to Swi4p (Iyer *et al.* 2001; MacIsaac *et al.* 2006). This regulatory element is near two Ste12p-binding sites ([A]TGAAACA) at 474-481 and 522-530 bp (Cullen *et al.* 2004). Ste12p regulates the *MSB2* promoter to induce its expression through positive feedback. Although Swi4/6p are positive regulators of G1/S transcription, the SBF complex is associated with the transcriptional repressor, Whi5p in early G1 (Costanzo *et al.* 2004; de Bruin *et al.* 2004; Palumbo *et al.* 2016). Therefore, the SBF complex might regulate *MSB2* expression in a number of ways. Interestingly, growth of cells in the non-preferred carbon source, GAL partially overrides the cell-cycle regulation of the fMAPK pathway. This might result from induction of *MSB2*expression by a starvation-dependent transcription factor, and/or by elevated processing of the Msb2p protein, which occurs at elevated levels in galactose (Vadaie *et al.* 2008).

We also show that the gene encoding the adaptor Sho1p is also subject to cell-cycle control. Several transcription factors that regulate cell-cycle progression bind the *SHO1* promoter, including Fhk1p (Ostrow *et al.* 2014) and Mbp1p (MacIsaac *et al.* 2006). Whether these proteins impart cell-cycle regulation of Sho1p levels remains to be determined. It has previously been shown that *TEC1* expression is induced at M/G1 boundary by Swi5p transcription factor (Cho *et al.* 1998; Spellman *et al.* 1998; Wittenberg and Reed 2005). Therefore, cell-cycle regulation of the fMAPK pathway may occur through multiple mechanisms.

Coupling the activity of the fMAPK pathway to the cell cycle may occur for the pathway to regulate intrinsic polarity, which occurs in G1, under some conditions (Prabhakar *et al.* 2020). Coupling the activity of the fMAPK pathway to the cell cycle may also impact its ability to regulate filamentous growth. We show here that the major cell adhesion molecule and flocculin Flo11p is regulated throughout the cell cycle. Flo11p is required to control adhesion functions under a variety of conditions, including nutrient-replete conditions (Pitoniak *et al.* 2009; Basu *et al.* 2016) and may have biological effects on mating (Guo *et al.* 2000). Intriguingly, another target of the MAPK pathway, *BUD8*(Adhikari and Cullen 2014), which marks the distal pole and is required for distal budding during filamentous growth (Taheri *et al.* 2000; Harkins *et al.* 2001; Cullen and Sprague 2002), is also cell-cycle regulated (Schenkman *et al.* 2002). The fMAPK pathway also regulates cell-cycle progression by controlling *CLN1* expression. Therefore the cell-cycle regulation of a differentiation-type MAPK pathway that itself alters the cell cycle might be critical for its morphogenetic responses to be coordinated. Interestingly, human MEK and ERK are also activated during mitosis in somatic cells to regulate the spindle assembly checkpoint (Shapiro *et al.* 1998; Horne and Guadagno 2003; Rosner 2007; Cao *et al.* 2010), proper entry into anaphase (Shapiro *et al.* 1998; Roberts *et al.* 2002), and fragmentation of Golgi cisternae (Acharya *et al.* 1998; Cha and Shapiro 2001; Shaul and Seger 2006). Therefore, MAP kinases may have a general role in regulating events that occur throughout the cell cycle (Pages *et al.* 1993; Mansour *et al.* 1994; Wright *et al.* 1999; Katz *et al.* 2007).

### The Mother-Bud Neck: A Hub for fMAPK Pathway Signaling?

Cumulatively, we have now amassed evidence that proteins that primarily function at the mother-bud neck regulate the fMAPK pathway. These include axial markers that control budsite-selection (Basu *et al.* 2016), cytokinesis remnant proteins (Prabhakar *et al.* 2020), the septins themselves (this study), and the cytokinesis regulator Hof1 (this study). A subset of these proteins may function to regulate the localization and/or activity of the adaptor protein Sho1p, which also localized to the mother-bud neck. Sho1p interacts with Hof1p and has functions in cytokinesis (Labedzka *et al.* 2012). Interestingly, the fMAP kinase Kss1p and cell wall integrity kinase Slt2p also act in septum assembly during cytokinesis (PÉrez *et al.* 2016). Thus, a fMAPK pathway complex may function at the neck to coordinate cytokinesis and next round of bud emergence. The spatial localization of these proteins can be viewed as a type of compartmentalization, which is a classic mode for maintaining signal specificity (Ebisuya *et al.* 2005; Doncic *et al.* 2015). Compartmentalization occurs on many levels, by restricting signaling to different cell types, organelles, parts of the plasma membrane, and even at different points in the cell cycle (Henis *et al.* 2009).

### A Positive Feedback Loop by One Pathway Sets up a Negative Feedback Loop by A Pathway with Shared Components

Most signaling pathways share components with other pathways. Pathway interactions can allow for modulation of pathway outputs. Here we provide evidence for cross-pathway feedback between the fMAPK pathway and the HOG pathway. Positive feedback through the fMAPK pathway induces HOG pathway activity, presumably to modulate fMAPK pathway activity. Interestingly, bleed through to the HOG pathway creates a loss of signal from the positive-feedback loop and a negative signal by stimulation of an antagonistic pathway. Such cross feedback might also impact target gene expression, although overexpression of Msb2p induces a non-overlapping set of targets as overexpression of Hkr1p (Pitoniak *et al.* 2009). The cross-pathway feedback also fits with the fact that the HOG pathway is induced by non-preferred carbon sources (Adhikari and Cullen 2014). Such modulation may fine-tune fMAPK pathway activity, which needs to be at the right level to promote filamentous growth and bud emergence, and at elevated levels, can lead to morphogenetic problems.

### General Ramifications to Pathway Specificity

Pathway specificity is one of the central questions in the signaling field. Signal duration, magnitude and subcellular compartmentalization of pathway regulators can have a profound impact on signal specificity and cellular outputs. It may be possible that the temporal regulation, by cell-cycle regulation of *MSB2* and *SHO1* gene expression, might be directly mechanistically connected to their spatial regulation, such as Sho1p’s localization at the mother-bud neck in M/G1. However, this need not necessarily be the case. Because both proteins are required to activate the fMAPK pathway, temporal and spatial control might acts in a coincidence manner to insure precise spatiotemporal activation of the fMAPK pathway. Although signaling pathways sense and respond to unique stimuli, often times multiple MAPK pathways collaborate to generate an appropriate response (Errede *et al.* 1995; Zarzov *et al.* 1996; Buehrer and Errede 1997; BaltanÁs *et al.* 2013; Adhikari and Cullen 2014; Prabhakar *et al.* 2020). Differences in activity between pathways that share components throughout the cell cycle may impact specificity and cell differentiation.

## MATERIALS AND METHODS

### Strains and Plasmids

Yeast strains are described in Table 1. Gene disruptions were made with antibiotic resistance markers *KanMX6*(Longtine *et al.* 1998), *HYG* and *NAT*(Goldstein and McCusker 1999) using PCR-based methods. Pop-in pop-out strategy was used to make internal epitope fusions (Schneider *et al.* 1995). Some strains were made *ura3-* by selection on 5-fluoroorotic acid (5-FOA). Gene disruptions were confirmed by PCR-based Southern analysis and also by phenotype when applicable. The *swi6*Δ mutant had a severe growth defect, which prevented evaluation of fMAPK and HOG pathway activities.

Most of the plasmids used in this study belong to the pRS series of plasmids (pRS315 and pRS316)(Sikorski and Hieter 1989). *pGFP-MSB2*(Adhikari *et al.* 2015b), *pRS316-SHO1-GFP*(Marles *et al.* 2004), p*SHO1*^P120L^(Vadaie *et al.* 2008), p*SHO1-GFP::NAT*(Prabhakar *et al.* 2020), pMyr-*SHO1* and pMyrAS-*SHO1*(Raitt *et al.* 2000), YCp50-*STE11-4*(Stevenson *et al.* 1992) and p*STE4*(Stevenson *et al.* 1992) have been described.

### Microbial Techniques

Standard methods were followed during yeast and bacterial strain manipulations (Sambrook 1989; Rose 1990). Budding pattern was determined as described (Cullen and Sprague 2002). The activity of the *FUS1-HIS3*(McCaffrey *et al.* 1987) growth reporter in cells lacking an intact mating pathway (*ste4*Δ) is dependent on components of the fMAPK pathway (Cullen *et al.* 2004) and was determined by growth of cells on media lacking histidine and supplemented with ATA (3-amino-1,2,4-triazole) for 3 d. Beta-galactosidase assays to assess the activity of the *FUS1-lacZ* reporter were performed as described (Cullen *et al.* 2000). The single-cell invasive growth assay (Cullen and Sprague 2000) and the plate-washing assay (Roberts and Fink1994) have been previously described.

### Immunoblot Analysis

Immunoblot analysis to detect phosphorylated MAP kinases has been described (Sabbagh *et al.* 2001; Lee and Dohlman 2008; Basu *et al.* 2016; Prabhakar *et al.* 2020). Proteins were precipitated from cell pellets stored at −80°C by trichloroacetic acid (TCA) and analyzed on 10% sodium dodecyl sulfate polyacrylamide gel electrophoresis (SDS-PAGE). Proteins were transferred to nitrocellulose membranes (Amersham™ Protran™ Premium 0.45 μm NC, GE Healthcare Life sciences, 10600003). For Msb2p-HA and Hkr1p-HA blots, 6% acrylamide gel was used.

ERK-type MAP kinases (P~Kss1p, P~Fus3p and P~Slt2p) were detected using α-p44/42 antibodies (Cell Signaling Technology, Danvers, MA, 4370 and 9101) at a 1:5,000 dilution. 9101 gave a stronger signal for P~Kss1p over P~Fus3p, while 4370 detected both proteins with similar strength. α-p38-type antibody at 1:5,000 dilution (Cell Signaling Technology, Danvers, MA 9211) was used to detect P~Hog1p. α-HA antibody at a 1:5,000 dilution (Roche Diagnostics, 12CA5) was used to detect Clb2p-HA, Msb2p-HA and Hkr1p-HA. Clb2p-Myc was detected using α-c-Myc antibody at 1:5,000 dilution (Santa Cruz Biotechnolog, Dallas, TX, 9E10) and Sho1p-GFP was detected using α-GFP antibody at 1:5,000 dilution (Roche Diagnostics, clones 7.1 and 13.1, 11814460001). α-Pgk1 antibody was used at a 1:5,000 dilution for total protein levels (Novex, 459250). For secondary antibodies, goat α-rabbit secondary IgG-HRP antibody was used at a 1:10,000 dilution (Jackson ImmunoResearch Laboratories, Inc., West Grove, PA, 111-035-144). Goat α-mouse secondary IgG-HRP antibody was used at a 1:5,000 dilution (BioRad Laboratories, Hercules, CA, 170-6516). Phospho-MAPK antibodies were incubated in 1X TBST (10 mM TRIS-HCl pH 8, 150 mM NaCl, 0.05% Tween 20) with 5% BSA. For all other antibodies, 1X TBST with 5% non-fat dried milk was used. Primary incubations were carried out for 16 h at 4°C. Secondary incubations were carried out for 1 h at 25°C.

### Cell Synchronization and Cell-Cycle Experiments

Cell synchronization by elutriation (Rosebrock 2017) was not feasible for cells of the ∑1278b background because cells fail to separate, even those lacking the adhesion molecule Flo11p (Vandermeulen and Cullen 2020). Cell synchronization experiments were performed as previously described (Breeden 1997; Prabhakar *et al.* 2020). Overnight cultures were resuspended in fresh media and grown to an optical density (O.D.) A_600_ of 0.2 at 30°C. Strains that required synthetic media (SD-URA) to maintain plasmid selection were harvested and resuspended in equal volume of YEPD and incubated for 1 h at 30°C prior to α-factor treatment. 10 ml aliquot was harvested as asynchronous culture. To arrest cells in G1,α factor was added to a final concentration of 5 μg/ml and the culture was incubated for 2 h at 30°C. 10 ml aliquots were harvested at 5 min, 30 min and 2 h during α-factor treatment. To arrest cells in S phase, hydroxyurea (HU) (MilliporeSigma, Burlington, MA, H8627) was added to a final concentration of 400 mM and incubated for 4 h. Arrested cells were washed twice with water (pre-warmed at 30°C) and resuspended in fresh YEPD or YEP-GAL media (pre-warmed at 30°C) to release cells into the cell cycle. 10 ml aliquots were harvested every 10 or 20 min and stored at −80°C.

### DIC and Fluorescence Microscopy

Differential-interference-contrast (DIC) and fluorescence microscopy using FiTC and TRiTC filter sets were performed using an Axioplan 2 fluorescent microscope (Zeiss, Oberkochen, Germany) with a Plan-Apochromat 100X/1.4 (oil) objective (N.A. 1.4) (cover slip 0.17) (Zeiss, Oberkochen, Germany). Digital images were obtained at multiple focal planes with the Axiocam MRm camera (Zeiss, Oberkochen, Germany) and Axiovision 4.4 software (Zeiss, Oberkochen, Germany). Adjustments to brightness and contrast were made in Adobe Photoshop (Adobe, San Jose, CA).

Time-lapse microscopy was performed on a Zeiss 710 confocal microscope (Zeiss, Oberkochen, Germany) equipped with a Plan-Apochromat 40x/1.4 Oil DIC M27 objective. For GFP, 488nm laser (496nm-548nm filter); for mCherry, 580 nm laser (589nm-708nm filter); for YFP, 517 nm laser (532-620 filter) and for CFP, 458 nm laser (462-532 filter) were used. For Sho1p-GFP time lapse, 9 z-stacks 1μm thick; for GAL-GFP-Msb2p time lapse 6 z-stacks 0.6 μm thick; and for Hof1p-CFP and Sho1p-YFP co-localization, 8 z-stacks 1.2 μm thick were captured at 10 min intervals.

Cells for time-lapse and co-localization studies were prepared as described in (Prabhakar *et al.* 2020). Cells were grown at 30°C for 16 h in SD-URA and diluted to < 0.1 O.D. 10 μL of diluted cells were placed under agarose pad (1%) prepared inside a 12 mm Nunc glass base dish (150680, Thermo Scientific, Waltham, MA). 100 μl of water was placed in the dish to prevent the agarose pad from drying and the petri dish was incubated at 30°C for 4 h prior to imaging.

### Indirect Immunofluorescence

Indirect immunofluorescence was performed as previously described with following modifications (Amberg *et al.* 2006; Schnell *et al.* 2012). Cells were grown to mid-log stage and 8% fresh paraformaldehyde (PFA) prepared in PBS (pH 7.4) was added directly to the culture (final concentration, 4%) for 10 min with shaking. Cells were harvested for 3 min at 350g and resuspended in KM solution (40 mM KPO_4_ pH 6.5, 500 μM MgCl_2_) containing 4% PFA for 1 h at 30°C with gentle shaking. Cells were washed twice with KM solution and once with KM solution containing 1.2 M sorbitol. After the last wash, cells were resuspended in 500 μl KM solution containing 1.2 M sorbitol and 60 μl Zymolyase (50 mg/ml 20T) for 20 min at 37°C. During zymolyase treatment, samples were periodically examined by DIC microscopy for cells with dull gray appearance and intact morphology (Niu *et al.* 2011). After Zymolyase treatment, cells were washed at 300g with KM solution containing 1.2 M sorbitol and resuspended in the same solution. Wells of Teflon-faced slides (MP Biomedicals, Santa Ana, CA, 096041205) were coated with 20 μl poly-L-lysine (Cultrex Poly-L-Lysine, Bio-Techne, Minneapolis, MN, 3438100-01) and incubated for 10 min at 24°C in a humid chamber. All solutions were centrifuged at 16,000g for 20 min at 4°C prior to adding to the wells. Wells were washed 5 times with 20 μl water and air dried. 20 μl of cells were spotted onto poly-L-lysine coated wells for 10 min at 24°C in the humid chamber. Excess solution was aspirated, and the slides were plunged into a coplin jar containing cold methanol for 6 minutes followed by cold acetone for 30 sec. Fixed and permeabilized cells were blocked using Image-iT™ FX Signal Enhancer (Thermo Fisher, Waltham, MA, i36933) for 30 min at 24°C in the humid chamber with gentle shaking. Excess solution was aspirated, and wells were washed 5 times with blocking buffer [PBS (pH 7.4), 5% normal goat serum (50062Z, Thermo Fisher, Waltham, MA), 1% BSA (80055-674, MilliporeSigma, Burlington, MA), 2% TritonX-100]. Wells were blocked again with 20 μl blocking buffer for 30 min at 24°C in the humid chamber with gentle shaking. After aspiration, cells were incubated with 20 μl of rabbit anti-p44/42 primary antibody (Cell Signaling Technology, Danvers, MA, 9101), prepared in blocking buffer at 1:20 dilution for 12 h at 24°C in a humid chamber with gentle shaking. Wells were washed 5 times for 5 min each with blocking buffer and co-stained with Goat anti-Rabbit IgG (H+L) Highly Cross-Adsorbed Secondary Antibody, Alexa Fluor 647 (Thermo Fisher, Waltham, MA, A-21245) and DAPI (4′,6-diamidino-2-phenylindole) at 1:1000 dilution each for 4 h at 24°C in a dark humid chamber with gentle shaking. Wells were washed 5 times for 10 min each with blocking buffer. After the last wash, wells were sealed with ProLong™ Diamond Antifade Mountant (Thermo Fisher, Waltham, MA, P36970) and covered with a cover slip. Slides were incubated in dark at 24°C for 24 h prior to imaging.

### Image Analysis

Quantitation of P~Kss1p by immunofluorescence is discussed elsewhere (Prabhakar and Cullen, *In Prep*) Briefly, total fluorescent intensity for each cell was measured in ImageJ by subtracting background intensity from the mean fluorescent intensity, which was recorded using *Analyze>Measure* option. Normalized fluorescent intensity for each cell was quantified as previously described (Okada *et al.* 2017; Prabhakar *et al.* 2020) using a custom MATLAB (MATLAB R2016b, The MathWorks, Inc., Natick, MA) code (Prabhakar *et al.* 2020). For each cell, pixel intensities greater than the mean+2 STD were background subtracted, normalized to the peak value (which was set to 1), and summed.

For time-lapse microscopy, raw images were imported into ImageJ. Cells were registered using HyperStackReg plugin (ThÉvenaz *et al.* 1998; Sharma 2018) to remove drift in the position of cells that occurred during imaging. Grayscale fluorescence images were converted to maximum intensity projection and inverted. Kymographs were performed as described (Prabhakar *et al.* 2020).

### Quantitative PCR Analysis

Quantitative RT-qPCR was performed as previously described (Adhikari and Cullen 2014; Chow *et al.* 2019a; Prabhakar *et al.* 2020). Samples harvested during cell-cycle experiments were used for total RNA extraction, which was done by hot acid phenol-chloroform treatment and further purified using RNeasy Mini Kit (Qiagen, Hilden, Germany, 74104). RNA stability was determined by agarose gel electrophoresis in 0.8% agarose Tris-Borato-EDTA (TBE, 89 mM Tris base, 89 mM Boric acid, 2mM EDTA). Concentration and purity were determined by absorbance using NanoDrop (NanoDrop, 2000C, Thermo Fisher Scientific, Waltham, MA). Concentration of total RNA was adjusted to 60 ng/μl and cDNA was synthesized using iScript Reverse Transcriptase Supermix (BioRad, Hercules, CA, 1708840). qPCR was performed using iTaq Universal SYBR Green Supermix (BioRad, Hercules, CA, 1725120) on BioRad thermocycler (CFX384 Real-Time System). Reactions contained 10 μl samples (2.5 μl 60 ng/μl cDNA, 0.2 μM each primer, 5 μl SYBRGreen master mix). Relative gene expression was calculated using the 2^−_Δ_Ct^ formula, where Ct is defined as the cycle at which fluorescence was determined to be statistically significant above background; ΔCt is the difference in Ct of the gene of interest and the housekeeping gene (*ACT1).* The primers used were: *MSB2* forward (5′-CACTGCAAGCAGGTGGCTCT-3′), *MSB2* reverse (5′-GAGGAGCCCGACAGTGTTGC-3′); *HKR1* forward (5′-AAACCATGGGCGAAAATGGC-3′), *HKR1* Reverse (5′-AAGGCAGGGGCTGTGAATAC-3′); *KSS1* forward (5′-CCCAAGTGATGAGCCGGAAT-3′), *KSS1* reverse (5′-TGGGCACTTCTTCCTCCTCT-3′); *SHO1* forward (5′-AACTACGATGGGAGACACTTTG-3′), *SHO1* reverse (5′-TCGTAAGCATCATCGTCATCAG-3′) (Adhikari and Cullen 2014); *TEC1* forward (5′-ATGTTTCCAGAAGCCGTAGTT-3′), *TEC1* reverse (5′-TTTAGCACCCAGTCCAGTATTT-3′) (Adhikari and Cullen 2014); *STE12* forward (5′-GCAATCTTACCCAAACGGAATG-3′), *STE12* reverse (5′-AATCGTCCGCGCCATAAA-3′) (Adhikari and Cullen 2014); *FLO11* forward (5′-CACTTTTGAAGTTTATGCCACACAAG-3′), *FLO11* reverse (5′-CTTGCATATTGAGCGGCACTAC-3′) (Chen and Fink 2006) and *ACT1* forward (5′-TGGATTCCGGTGATGGTGTT-3′), *ACT1* reverse (5′-CGGCCAAATCGATTCTCAA-3′) (Chow *et al.* 2019b). Experiments were performed with two independent biological replicates and two technical replicates for each biological replicate.

### Statistical Analysis

Statistical tests and sample size (n) have been described in figure legends wherever applicable. Statistical analyses were performed in Microsoft Excel and Minitab (www.minitab.com). Oneway ANOVA with Tukey’s test and/or Dunnett’s test was used for statistical analysis.

## ABBREVIATIONS

ATA: (3-amino-1,2,4-triazole)
5-FOA: 5-fluoroorotic acid
CFP: cyan fluorescent protein
D: dextrose
DAPI: 4′,6-diamidino-2-phenylindole
DIC: differential interference contrast
GAL: galactose
GAP: GTPase activating protein
GEF: guanine nucleotide exchange factor
GTPase: guanine nucleotide triphosphatase
GFP: green fluorescent protein
GLU: glucose
GAL: galactose
HA: hemaglutinin
HOG: high osmolarity glycerol response
HU: hydroxyurea
MAPK: mitogen activated protein kinase
O.D.: optical density
PAK: p21 activated kinase
PFA: parafolmaldehyde
RT-qPCR: Reverse transcriptase quantitative polymerase chain reaction
PM: plasma membrane
Rho: Ras homology
SDS-PAGE: sodium dodecyl sulfatepolyacrylamide gel electrophoresis
S.E.M: standard error of mean
TBE: Tris-Borate-EDTA
TCA: trichloroacetic acid
WT: wild type
YFP: yellow fluorescent protein
YNB: Yeast Nitrogen base.

## ACKNOWLEDGEMENTS

The work was supported from a grant from the NIH (GM#098629). Thanks to Haruo Saito (University of Toyko) for reagents. Andrew Pitoniak helped with experiments.

## REFERENCES

Acharya, U., A. Mallabiabarrena, J. K. Acharya and V. Malhotra, 1998 Signaling via mitogen-activated protein kinase kinase (MEK1) is required for Golgi fragmentation during mitosis. Cell 92:183–192.

Adhikari, H., L. M. Caccamise, T. Pande and P. J. Cullen, 2015a Comparative Analysis of Transmembrane Regulators of the Filamentous Growth Mitogen-Activated Protein Kinase Pathway Uncovers Functional and Regulatory Differences. Eukaryot Cell 14:868–883.

Adhikari, H., and P. J. Cullen, 2014 Metabolic respiration induces AMPK-and Ire1p-dependent activation of the p38-Type HOG MAPK pathway. PLoS Genet 10:e1004734.

Adhikari, H., N. Vadaie, J. Chow, L. M. Caccamise, C. A. Chavel et al., 2015b Role of the unfolded protein response in regulating the mucin-dependent filamentous-growth mitogen-activated protein kinase pathway. Mol Cell Biol 35:1414–1432.

Ahn, S. H., A. Acurio and S. J. Kron, 1999 Regulation of G2/M progression by the STE mitogen-activated protein kinase pathway in budding yeast filamentous growth. Mol Biol Cell 10:3301–3316.

Aktas, H., H. Cai and G. M. Cooper, 1997 Ras links growth factor signaling to the cell cycle machinery via regulation of cyclin D1 and the Cdk inhibitor p27KIP1. Mol Cell Biol 17:3850–3857.

Amberg, D. C., D. J. Burke and J. N. Strathern, 2006 Yeast Immunofluorescence. Cold Spring Harbor Protocols 2006:pdb.prot4167.

Andrews, B. J., and I. Herskowitz, 1989 The yeast SWI4 protein contains a motif present in developmental regulators and is part of a complex involved in cell-cycle-dependent transcription. Nature 342:830–833.

Baltanás, R., A. Bush, A. Couto, L. Durrieu, S. Hohmann et al., 2013 Pheromone-induced morphogenesis improves osmoadaptation capacity by activating the HOG MAPK pathway. Sci Signal 6:ra26.

Bao, M. Z., M. A. Schwartz, G. T. Cantin, J. R. Yates, 3rd and H. D. Madhani, 2004 Pheromone-dependent destruction of the Tec1 transcription factor is required for MAP kinase signaling specificity in yeast. Cell 119:991–1000.

Bardwell, L., 2004 A walk-through of the yeast mating pheromone response pathway. Peptides 25:1465–1476.

Bardwell, L., 2006 Mechanisms of MAPK signalling specificity. Biochem Soc Trans 34:837–841.

Bardwell, L., J. G. Cook, E. C. Chang, B. R. Cairns and J. Thorner, 1996 Signaling in the yeast pheromone response pathway: specific and high-affinity interaction of the mitogen-activated protein (MAP) kinases Kss1 and Fus3 with the upstream MAP kinase kinase Ste7. Mol Cell Biol 16:3637–3650.

Bardwell, L., J. G. Cook, D. Voora, D. M. Baggott, A. R. Martinez et al., 1998 Repression of yeast Ste12 transcription factor by direct binding of unphosphorylated Kss1 MAPK and its regulation by the Ste7 MEK. Genes Dev 12:2887–2898.

Basu, S., B. González, B. Li, G. Kimble, K. G. Kozminski et al., 2020 Functions for Cdc42p BEM adaptors in regulating a differentiation-type MAP kinase pathway. Mol Biol Cell 31:491–510.

Basu, S., N. Vadaie, A. Prabhakar, B. Li, H. Adhikari et al., 2016 Spatial landmarks regulate a Cdc42-dependent MAPK pathway to control differentiation and the response to positional compromise. Proc Natl Acad Sci U S A 113:E2019–2028.

Bi, E., and H. O. Park, 2012 Cell polarization and cytokinesis in budding yeast. Genetics 191:347–387.

Birkaya, B., A. Maddi, J. Joshi, S. J. Free and P. J. Cullen, 2009 Role of the cell wall integrity and filamentous growth mitogen-activated protein kinase pathways in cell wall remodeling during filamentous growth. Eukaryot Cell 8:1118–1133.

Breeden, L., and K. Nasmyth, 1987 Cell cycle control of the yeast HO gene: cis-and trans-acting regulators. Cell 48:389–397.

Breeden, L. L., 1997 α-Factor synchronization of budding yeast, pp. 332–342 in Methods in Enzymology. Academic Press.

Breitkreutz, A., and M. Tyers, 2002 MAPK signaling specificity: it takes two to tango. Trends Cell Biol 12:254–257.

Breker, M., M. Gymrek and M. Schuldiner, 2013 A novel single-cell screening platform reveals proteome plasticity during yeast stress responses. J Cell Biol 200:839–850.

Brewster, J. L., T. de Valoir, N. D. Dwyer, E. Winter and M. C. Gustin, 1993 An osmosensing signal transduction pathway in yeast. Science 259:1760–1763.

Brückner, S., T. Köhler, G. H. Braus, B. Heise, M. Bolte et al., 2004 Differential regulation of Tec1 by Fus3 and Kss1 confers signaling specificity in yeast development. Curr Genet 46:331–342.

Buehrer, B. M., and B. Errede, 1997 Coordination of the mating and cell integrity mitogen-activated protein kinase pathways in Saccharomyces cerevisiae. Mol Cell Biol 17:6517–6525.

Cao, J. N., N. Shafee, L. Vickery, S. Kaluz, N. Ru et al., 2010 Mitogen-activated protein/extracellular signal-regulated kinase kinase 1act/tubulin interaction is an important determinant of mitotic stability in cultured HT1080 human fibrosarcoma cells. Cancer Res 70:6004–6014.

Carlson, M., and D. Botstein, 1982 Two differentially regulated mRNAs with different 5′ ends encode secreted with intracellular forms of yeast invertase. Cell 28:145–154.

Cepeda-Garcia, C., 2017 Determination of Cell Cycle Stage and Mitotic Exit Through the Quantification of the Protein Levels of Known Mitotic Regulators. Methods Mol Biol 1505:45–57.

Cha, H., and P. Shapiro, 2001 Tyrosine-phosphorylated extracellular signal--regulated kinase associates with the Golgi complex during G2/M phase of the cell cycle: evidence for regulation of Golgi structure. J Cell Biol 153:1355–1367.

Chant, J., M. Mischke, E. Mitchell, I. Herskowitz and J. R. Pringle, 1995 Role of Bud3p in producing the axial budding pattern of yeast. J Cell Biol 129:767–778.

Chavel, C. A., H. M. Dionne, B. Birkaya, J. Joshi and P. J. Cullen, 2010 Multiple Signals Converge on a Differentiation MAPK Pathway. PLOS Genetics 6:e1000883.

Chen, H., and G. R. Fink, 2006 Feedback control of morphogenesis in fungi by aromatic alcohols. Genes & Development 20:1150–1161.

Chen, R. E., and J. Thorner, 2007 Function and regulation in MAPK signaling pathways: lessons learned from the yeast Saccharomyces cerevisiae. Biochim Biophys Acta 1773:1311–1340.

Chen, R. H., C. Sarnecki and J. Blenis, 1992 Nuclear localization and regulation of erk-and rsk-encoded protein kinases. Mol Cell Biol 12:915–927.

Cho, R. J., M. J. Campbell, E. A. Winzeler, L. Steinmetz, A. Conway et al., 1998 A genome-wide transcriptional analysis of the mitotic cell cycle. Mol Cell 2:65–73.

Chou, S., L. Huang and H. Liu, 2004 Fus3-regulated Tec1 degradation through SCFCdc4 determines MAPK signaling specificity during mating in yeast. Cell 119:981–990.

Chow, J., H. M. Dionne, A. Prabhakar, A. Mehrotra, J. Somboonthum et al., 2019a Aggregate Filamentous Growth Responses in Yeast. mSphere 4:e00702–00718.

Chow, J., I. Starr, S. Jamalzadeh, O. Muniz, A. Kumar et al., 2019b Filamentation Regulatory Pathways Control Adhesion-Dependent Surface Responses in Yeast. Genetics 212:667690.

Cook, J. G., L. Bardwell and J. Thorner, 1997 Inhibitory and activating functions for MAPK Kss1 in the S. cerevisiae filamentous-growth signalling pathway. Nature 390:85–88.

Costanzo, M., J. L. Nishikawa, X. Tang, J. S. Millman, O. Schub et al., 2004 CDK activity antagonizes Whi5, an inhibitor of G1/S transcription in yeast. Cell 117:899–913.

Cross, F. R., L. Schroeder, M. Kruse and K. C. Chen, 2005 Quantitative characterization of a mitotic cyclin threshold regulating exit from mitosis. Molecular biology of the cell 16:2129–2138.

Cullen, P. J., W. Sabbagh, Jr., E. Graham, M. M. Irick, E. K. van Olden et al., 2004 A signaling mucin at the head of the Cdc42-and MAPK-dependent filamentous growth pathway in yeast. Genes Dev 18:1695–1708.

Cullen, P. J., J. Schultz, J. Horecka, B. J. Stevenson, Y. Jigami et al., 2000 Defects in protein glycosylation cause SHO1-dependent activation of a STE12 signaling pathway in yeast. Genetics 155:1005–1018.

Cullen, P. J., and G. F. Sprague, Jr., 2000 Glucose depletion causes haploid invasive growth in yeast. Proc Natl Acad Sci U S A 97:13619–13624.

Cullen, P. J., and G. F. Sprague, Jr., 2002 The roles of bud-site-selection proteins during haploid invasive growth in yeast. Mol Biol Cell 13:2990–3004.

Dangi, S., F. M. Chen and P. Shapiro, 2006 Activation of extracellular signal-regulated kinase (ERK) in G2 phase delays mitotic entry through p21CIP1. Cell Prolif 39:261–279.

Davenport, K. D., K. E. Williams, B. D. Ullmann and M. C. Gustin, 1999 Activation of the Saccharomyces cerevisiae filamentation/invasion pathway by osmotic stress in high-osmolarity glycogen pathway mutants. Genetics 153:1091–1103.

de Bruin, R. A., W. H. McDonald, T. I. Kalashnikova, J. Yates, 3rd and C. Wittenberg, 2004 Cln3 activates G1-specific transcription via phosphorylation of the SBF bound repressor Whi5. Cell 117:887–898.

De Vit, M. J., J. A. Waddle and M. Johnston, 1997 Regulated nuclear translocation of the Mig1 glucose repressor. Mol Biol Cell 8:1603–1618.

Dinsmore, C. J., and P. Soriano, 2018 MAPK and PI3K signaling: At the crossroads of neural crest development. Dev Biol 444 Suppl 1:S79–s97.

Doncic, A., O. Atay, E. Valk, A. Grande, A. Bush et al., 2015 Compartmentalization of a bistable switch enables memory to cross a feedback-driven transition. Cell 160:1182–1195.

Ebisuya, M., K. Kondoh and E. Nishida, 2005 The duration, magnitude and compartmentalization of ERK MAP kinase activity: mechanisms for providing signaling specificity. J Cell Sci 118:2997–3002.

Elion, E. A., B. Satterberg and J. E. Kranz, 1993 FUS3 phosphorylates multiple components of the mating signal transduction cascade: evidence for STE12 and FAR1. Mol Biol Cell 4:495–510.

Eluere, R., N. Offner, I. Varlet, O. Motteux, L. Signon et al., 2007 Compartmentalization of the functions and regulation of the mitotic cyclin Clb2 in S. cerevisiae. J Cell Sci 120:702–711.

Errede, B., R. M. Cade, B. M. Yashar, Y. Kamada, D. E. Levin et al., 1995 Dynamics and organization of MAP kinase signal pathways. Mol Reprod Dev 42:477–485.

Fischer, M. S., V. W. Wu, J. E. Lee, R. C. O’Malley and N. L. Glass, 2018 Regulation of Cell-to-Cell Communication and Cell Wall Integrity by a Network of MAP Kinase Pathways and Transcription Factors in Neurospora crassa. Genetics 209:489–506.

Gancedo, J. M., 2001 Control of pseudohyphae formation in Saccharomyces cerevisiae. FEMS Microbiol Rev 25:107–123.

Gimeno, C. J., P. O. Ljungdahl, C. A. Styles and G. R. Fink, 1992 Unipolar cell divisions in the yeast S. cerevisiae lead to filamentous growth: regulation by starvation and RAS. Cell 68:1077–1090.

Goldstein, A. L., and J. H. McCusker, 1999 Three new dominant drug resistance cassettes for gene disruption in Saccharomyces cerevisiae. Yeast 15:1541–1553.

Guo, B., C. A. Styles, Q. Feng and G. R. Fink, 2000 A Saccharomyces gene family involved in invasive growth, cell-cell adhesion, and mating. Proc Natl Acad Sci U S A 97:12158–12163.

Hall, P. A., Hilary Russell, S.E., and J.R. Pringle, 2008 The Septins. Wiley-Blackwell.

Hao, N., Y. Zeng, T. C. Elston and H. G. Dohlman, 2008 Control of MAPK specificity by feedback phosphorylation of shared adaptor protein Ste50. J Biol Chem 283:33798–33802.

Harkins, H. A., N. Pagé, L. R. Schenkman, C. De Virgilio, S. Shaw et al., 2001 Bud8p and Bud9p, proteins that may mark the sites for bipolar budding in yeast. Mol Biol Cell 12:2497–2518.

Henis, Y. I., J. F. Hancock and I. A. Prior, 2009 Ras acylation, compartmentalization and signaling nanoclusters (Review). Mol Membr Biol 26:80–92.

Hirosumi, J., G. Tuncman, L. Chang, C. Z. Görgün, K. T. Uysal et al., 2002 A central role for JNK in obesity and insulin resistance. Nature 420:333–336.

Horne, M. M., and T. M. Guadagno, 2003 A requirement for MAP kinase in the assembly and maintenance of the mitotic spindle. J Cell Biol 161:1021–1028.

Ingram, A. J., L. James, L. Cai, K. Thai, H. Ly et al., 2000 NO inhibits stretch-induced MAPK activity by cytoskeletal disruption. J Biol Chem 275:40301–40306.

Irniger, S., S. Piatti, C. Michaelis and K. Nasmyth, 1995 Genes involved in sister chromatid separation are needed for B-type cyclin proteolysis in budding yeast. Cell 81:269–278.

Iyer, V. R., C. E. Horak, C. S. Scafe, D. Botstein, M. Snyder et al., 2001 Genomic binding sites of the yeast cell-cycle transcription factors SBF and MBF. Nature 409:533–538.

Johnson, D. I., 1999 Cdc42: An essential Rho-type GTPase controlling eukaryotic cell polarity. Microbiol Mol Biol Rev 63:54–105.

Johnson, G. L., and R. Lapadat, 2002 Mitogen-activated protein kinase pathways mediated by ERK, JNK, and p38 protein kinases. Science 298:1911–1912.

Jordan, J. D., E. M. Landau and R. Iyengar, 2000 Signaling networks: the origins of cellular multitasking. Cell 103:193–200.

Karunanithi, S., and P. J. Cullen, 2012 The Filamentous Growth MAPK Pathway Responds to Glucose Starvation Through the Mig1/2 Transcriptional Repressors in Saccharomyces cerevisiae. Genetics 192:869–887.

Katz, M., I. Amit and Y. Yarden, 2007 Regulation of MAPKs by growth factors and receptor tyrosine kinases. Biochim Biophys Acta 1773:1161–1176.

Kim, H. B., B. K. Haarer and J. R. Pringle, 1991 Cellular morphogenesis in the Saccharomyces cerevisiae cell cycle: localization of the CDC3 gene product and the timing of events at the budding site. J Cell Biol 112:535–544.

Koç, A., L. J. Wheeler, C. K. Mathews and G. F. Merrill, 2004 Hydroxyurea arrests DNA replication by a mechanism that preserves basal dNTP pools. J Biol Chem 279:223–230.

Köhler, T., S. Wesche, N. Taheri, G. H. Braus and H. U. Mösch, 2002 Dual role of the Saccharomyces cerevisiae TEA/ATTS family transcription factor Tec1p in regulation of gene expression and cellular development. Eukaryot Cell 1:673–686.

Kuczera, T., O. Bayram, F. Sari, G. H. Braus and S. Irniger, 2010 Dissection of mitotic functions of the yeast cyclin Clb2. Cell Cycle 9:2611–2619.

Labedzka, K., C. Tian, U. Nussbaumer, S. Timmermann, P. Walther et al., 2012 Sho1p connects the plasma membrane with proteins of the cytokinesis network through multiple isomeric interaction states. J Cell Sci 125:4103–4113.

Lavoie, H., J. Gagnon and M. Therrien, 2020 ERK signalling: a master regulator of cell behaviour, life and fate. Nature Reviews Molecular Cell Biology 21:607–632.

Lawrence, M. C., A. Jivan, C. Shao, L. Duan, D. Goad et al., 2008 The roles of MAPKs in disease. Cell Research 18:436–442.

Leberer, E., C. Wu, T. Leeuw, A. Fourest-Lieuvin, J. E. Segall et al., 1997 Functional characterization of the Cdc42p binding domain of yeast Ste20p protein kinase. Embo j 16:83–97.

Lee, J. C., S. Kumar, D. E. Griswold, D. C. Underwood, B. J. Votta et al., 2000 Inhibition of p38 MAP kinase as a therapeutic strategy. Immunopharmacology 47:185–201.

Lee, K. S., K. Irie, Y. Gotoh, Y. Watanabe, H. Araki et al., 1993 A yeast mitogen-activated protein kinase homolog (Mpk1p) mediates signalling by protein kinase C. Mol Cell Biol 13:3067–3075.

Lee, M. J., and H. G. Dohlman, 2008 Coactivation of G protein signaling by cell-surface receptors and an intracellular exchange factor. Curr Biol 18:211–215.

Lenormand, P., C. Sardet, G. Pagès, G. L’Allemain, A. Brunet et al., 1993 Growth factors induce nuclear translocation of MAP kinases (p42mapk and p44mapk) but not of their activator MAP kinase kinase (p45mapkk) in fibroblasts. J Cell Biol 122:1079–1088.

Leone, G., J. DeGregori, R. Sears, L. Jakoi and J. R. Nevins, 1997 Myc and Ras collaborate in inducing accumulation of active cyclin E/Cdk2 and E2F. Nature 387:422–426.

Lippincott, J., K. B. Shannon, W. Shou, R. J. Deshaies and R. Li, 2001 The Tem1 small GTPase controls actomyosin and septin dynamics during cytokinesis. J Cell Sci 114:1379–1386.

Liu, H., C. A. Styles and G. R. Fink, 1993 Elements of the yeast pheromone response pathway required for filamentous growth of diploids. Science 262:1741–1744.

Lo, H. J., J. R. Kohler, B. DiDomenico, D. Loebenberg, A. Cacciapuoti et al., 1997 Nonfilamentous C. albicans mutants are avirulent. Cell 90:939–949.

Loeb, J. D., T. A. Kerentseva, T. Pan, M. Sepulveda-Becerra and H. Liu, 1999 Saccharomyces cerevisiae G1 cyclins are differentially involved in invasive and pseudohyphal growth independent of the filamentation mitogen-activated protein kinase pathway. Genetics 153:1535–1546.

Longtine, M. S., A. McKenzie, 3rd, D. J. Demarini, N. G. Shah, A. Wach et al., 1998 Additional modules for versatile and economical PCR-based gene deletion and modification in Saccharomyces cerevisiae. Yeast 14:953–961.

Ma, D., J. G. Cook and J. Thorner, 1995 Phosphorylation and localization of Kss1, a MAP kinase of the Saccharomyces cerevisiae pheromone response pathway. Mol Biol Cell 6:889–909.

MacIsaac, K. D., T. Wang, D. B. Gordon, D. K. Gifford, G. D. Stormo et al., 2006 An improved map of conserved regulatory sites for Saccharomyces cerevisiae. BMC Bioinformatics 7:113.

Madhani, H. D., and G. R. Fink, 1997 Combinatorial control required for the specificity of yeast MAPK signaling. Science 275:1314–1317.

Madhani, H. D., T. Galitski, E. S. Lander and G. R. Fink, 1999 Effectors of a developmental mitogen-activated protein kinase cascade revealed by expression signatures of signaling mutants. Proc Natl Acad Sci U S A 96:12530–12535.

Madhani, H. D., C. A. Styles and G. R. Fink, 1997 MAP kinases with distinct inhibitory functions impart signaling specificity during yeast differentiation. Cell 91:673–684.

Maeda, T., S. M. Wurgler-Murphy and H. Saito, 1994 A two-component system that regulates an osmosensing MAP kinase cascade in yeast. Nature 369:242–245.

Maekawa, M., T. Yamamoto, T. Tanoue, Y. Yuasa, O. Chisaka et al., 2005 Requirement of the MAP kinase signaling pathways for mouse preimplantation development. Development 132:1773–1783.

Mansour, S. J., W. T. Matten, A. S. Hermann, J. M. Candia, S. Rong et al., 1994 Transformation of mammalian cells by constitutively active MAP kinase kinase. Science 265:966–970.

Marles, J. A., S. Dahesh, J. Haynes, B. J. Andrews and A. R. Davidson, 2004 Protein-protein interaction affinity plays a crucial role in controlling the Sho1p-mediated signal transduction pathway in yeast. Mol Cell 14:813–823.

McCaffrey, G., F. J. Clay, K. Kelsay and G. F. Sprague, Jr., 1987 Identification and regulation of a gene required for cell fusion during mating of the yeast Saccharomyces cerevisiae. Mol Cell Biol 7:2680–2690.

McMurray, M. A., and J. Thorner, 2009 Septins: molecular partitioning and the generation of cellular asymmetry. Cell Div 4:18.

Meitinger, F., M. E. Boehm, A. Hofmann, B. Hub, H. Zentgraf et al., 2011 Phosphorylation-dependent regulation of the F-BAR protein Hof1 during cytokinesis. Genes Dev 25:875–888.

Meloche, S., and J. Pouysségur, 2007 The ERK1/2 mitogen-activated protein kinase pathway as a master regulator of the G1-to S-phase transition. Oncogene 26:3227–3239.

Molgaard, S., M. Ulrichsen, D. Olsen and S. Glerup, 2016 Detection of phosphorylated Akt and MAPK in cell culture assays. MethodsX 3:386–398.

Mosch, H. U., R. L. Roberts and G. R. Fink, 1996 Ras2 signals via the Cdc42/Ste20/mitogen-activated protein kinase module to induce filamentous growth in Saccharomyces cerevisiae. Proc Natl Acad Sci U S A 93:5352–5356.

Murphy, L. O., S. Smith, R. H. Chen, D. C. Fingar and J. Blenis, 2002 Molecular interpretation of ERK signal duration by immediate early gene products. Nat Cell Biol 4:556–564.

Nasmyth, K., and L. Dirick, 1991 The role of SWI4 and SWI6 in the activity of G1 cyclins in yeast. Cell 66:995–1013.

Nehlin, J. O., M. Carlberg and H. Ronne, 1991 Control of yeast GAL genes by MIG1 repressor: a transcriptional cascade in the glucose response. Embo J 10:3373–3377.

Nelson, B., A. B. Parsons, M. Evangelista, K. Schaefer, K. Kennedy et al., 2004 Fus1p interacts with components of the Hog1p mitogen-activated protein kinase and Cdc42p morphogenesis signaling pathways to control cell fusion during yeast mating. Genetics 166:67–77.

Niu, W., G. T. Hart and E. M. Marcotte, 2011 High-throughput immunofluorescence microscopy using yeast spheroplast cell-based microarrays. Methods Mol Biol 706:83–95.

O’Rourke, S. M., and I. Herskowitz, 1998 The Hog1 MAPK prevents cross talk between the HOG and pheromone response MAPK pathways in Saccharomyces cerevisiae. Genes Dev 12:2874–2886.

O’Rourke, S. M., and I. Herskowitz, 2002 A third osmosensing branch in Saccharomyces cerevisiae requires the Msb2 protein and functions in parallel with the Sho1 branch. Mol Cell Biol 22:4739–4749.

Oh, Y., J. Schreiter, R. Nishihama, C. Wloka and E. Bi, 2013 Targeting and functional mechanisms of the cytokinesis-related F-BAR protein Hof1 during the cell cycle. Mol Biol Cell 24:1305–1320.

Okada, S., M. E. Lee, E. Bi and H.-O. Park, 2017 Probing Cdc42 Polarization Dynamics in Budding Yeast Using a Biosensor. Methods in enzymology 589:171–190.

Omori, S., M. Hida, H. Fujita, H. Takahashi, S. Tanimura et al., 2006 Extracellular signal-regulated kinase inhibition slows disease progression in mice with polycystic kidney disease. J Am Soc Nephrol 17:1604–1614.

Ostrow, A. Z., T. Nellimoottil, S. R. Knott, C. A. Fox, S. Tavaré et al., 2014 Fkh1 and Fkh2 bind multiple chromosomal elements in the S. cerevisiae genome with distinct specificities and cell cycle dynamics. PLoS one 9:e87647.

Pages, G., P. Lenormand, G. L’Allemain, J. C. Chambard, S. Meloche et al., 1993 Mitogen-activated protein kinases p42mapk and p44mapk are required for fibroblast proliferation. Proc Natl Acad Sci U S A 90:8319–8323.

Palumbo, P., M. Vanoni, V. Cusimano, S. Busti, F. Marano et al., 2016 Whi5 phosphorylation embedded in the G1/S network dynamically controls critical cell size and cell fate. Nat Commun 7:11372.

Pan, X., T. Harashima and J. Heitman, 2000 Signal transduction cascades regulating pseudohyphal differentiation of Saccharomyces cerevisiae. Curr opin Microbiol 3:567–572.

Papa, S., P. M. Choy and C. Bubici, 2019 The ERK and JNK pathways in the regulation of metabolic reprogramming. Oncogene 38:2223–2240.

Pelet, S., 2017 Nuclear relocation of Kss1 contributes to the specificity of the mating response. Scientific Reports 7:43636.

Pérez, J., I. Arcones, A. Gómez, V. Casquero and C. Roncero, 2016 Phosphorylation of Bni4 by MAP kinases contributes to septum assembly during yeast cytokinesis. FEMS Yeast Research 16.

Peter, M., A. Gartner, J. Horecka, G. Ammerer and I. Herskowitz, 1993 FAR1 links the signal transduction pathway to the cell cycle machinery in yeast. Cell 73:747–760.

Peter, M., and I. Herskowitz, 1994 Direct inhibition of the yeast cyclin-dependent kinase Cdc28-Cln by Far1. Science 265:1228–1231.

Peter, M., A. M. Neiman, H. O. Park, M. van Lohuizen and I. Herskowitz, 1996 Functional analysis of the interaction between the small GTP binding protein Cdc42 and the Ste20 protein kinase in yeast. Embo j 15:7046–7059.

Pitoniak, A., B. Birkaya, H. M. Dionne, N. Vadaie and P. J. Cullen, 2009 The Signaling Mucins Msb2 and Hkr1 Differentially Regulate the Filamentation Mitogen-activated Protein Kinase Pathway and Contribute to a Multimodal Response. Molecular Biology of the Cell 20:3101–3114.

Pitoniak, A., C. A. Chavel, J. Chow, J. Smith, D. Camara et al., 2015 Cdc42p-Interacting Protein Bem4p Regulates the Filamentous-Growth Mitogen-Activated Protein Kinase Pathway. Molecular and Cellular Biology 35:417–436.

Posas, F., and H. Saito, 1997 Osmotic activation of the HOG MAPK pathway via Ste11p MAPKKK: scaffold role of Pbs2p MAPKK. Science 276:1702–1705.

Pouysségur, J., V. Volmat and P. Lenormand, 2002 Fidelity and spatio-temporal control in MAP kinase (ERKs) signalling. Biochem Pharmacol 64:755–763.

Prabhakar, A., J. Chow, A. J. Siegel and P. J. Cullen, 2020 Regulation of intrinsic polarity establishment by a differentiation-type MAPK pathway. J Cell Sci.

Prabhakar, A., N. Vadaie, T. Krzystek and P. J. Cullen, 2019 Proteins That Interact with the Mucin-Type Glycoprotein Msb2p Include a Regulator of the Actin Cytoskeleton. Biochemistry 58:4842–4856.

Radmaneshfar, E., D. Kaloriti, M. C. Gustin, N. A. Gow, A. J. Brown et al., 2013 From START to FINISH: the influence of osmotic stress on the cell cycle. PLoS One 8:e68067.

Raitt, D. C., F. Posas and H. Saito, 2000 Yeast Cdc42 GTPase and Ste20 PAK-like kinase regulate Sho1-dependent activation of the Hog1 MAPK pathway. Embo j 19:4623–4631.

Raman, M., W. Chen and M. H. Cobb, 2007 Differential regulation and properties of MAPKs. Oncogene 26:3100–3112.

Reiser, V., D. C. Raitt and H. Saito, 2003 Yeast osmosensor Sln1 and plant cytokinin receptor Cre1 respond to changes in turgor pressure. J Cell Biol 161:1035–1040.

Richardson, H., D. J. Lew, M. Henze, K. Sugimoto and S. I. Reed, 1992 Cyclin-B homologs in Saccharomyces cerevisiae function in S phase and in G2. Genes Dev 6:2021–2034.

Roberts, C. J., B. Nelson, M. J. Marton, R. Stoughton, M. R. Meyer et al., 2000 Signaling and circuitry of multiple MAPK pathways revealed by a matrix of global gene expression profiles. Science 287:873–880.

Roberts, E. C., P. S. Shapiro, T. S. Nahreini, G. Pages, J. Pouyssegur et al., 2002 Distinct cell cycle timing requirements for extracellular signal-regulated kinase and phosphoinositide 3-kinase signaling pathways in somatic cell mitosis. Mol Cell Biol 22:7226–7241.

Roberts, P. J., and C. J. Der, 2007 Targeting the Raf-MEK-ERK mitogen-activated protein kinase cascade for the treatment of cancer. Oncogene 26:3291–3310.

Roberts, R. L., and G. R. Fink, 1994 Elements of a single MAP kinase cascade in Saccharomyces cerevisiae mediate two developmental programs in the same cell type: mating and invasive growth. Genes Dev 8:2974–2985.

Rodriguez-Viciana, P., O. Tetsu, W. E. Tidyman, A. L. Estep, B. A. Conger et al., 2006 Germline mutations in genes within the MAPK pathway cause cardio-facio-cutaneous syndrome. Science 311:1287–1290.

Rose, M. D., Winston, F., and Hieter, P., 1990 Methods in yeast genetics. Cold Spring Harbor Laboratory Press, Cold Spring Harbor, NY.

Rosebrock, A. P., 2017 Synchronization of Budding Yeast by Centrifugal Elutriation. Cold Spring Harb Protoc 2017.

Rosner, M. R., 2007 MAP kinase meets mitosis: a role for Raf Kinase Inhibitory Protein in spindle checkpoint regulation. Cell Div 2:1.

Roux, P. P., and J. Blenis, 2004 ERK and p38 MAPK-activated protein kinases: a family of protein kinases with diverse biological functions. Microbiol Mol Biol Rev 68:320–344.

Rua, D., B. T. Tobe and S. J. Kron, 2001 Cell cycle control of yeast filamentous growth. Curr opin Microbiol 4:720–727.

Rupp, S., E. Summers, H. J. Lo, H. Madhani and G. Fink, 1999 MAP kinase and cAMP filamentation signaling pathways converge on the unusually large promoter of the yeast FLO11 gene. Embo J 18:1257–1269.

Sabbagh, W., L. J. Flatauer, A. J. Bardwell and L. Bardwell, 2001 Specificity of MAP Kinase Signaling in Yeast Differentiation Involves Transient versus Sustained MAPK Activation. Molecular Cell 8:683–691.

Sambrook, J., Fritsch, E.F., and Maniatis, T., 1989 Molecular cloning: a laboratory manual. Cold Spring Harbor Laboratory Press, Cold Spring Harbor, NY.

Sanders, S. L., and I. Herskowitz, 1996 The BUD4 protein of yeast, required for axial budding, is localized to the mother/BUD neck in a cell cycle-dependent manner. J Cell Biol 134:413–427.

Schenkman, L. R., C. Caruso, N. Pagé and J. R. Pringle, 2002 The role of cell cycle-regulated expression in the localization of spatial landmark proteins in yeast. J Cell Biol 156:829–841.

Schneider, B. L., W. Seufert, B. Steiner, Q. H. Yang and A. B. Futcher, 1995 Use of polymerase chain reaction epitope tagging for protein tagging in Saccharomyces cerevisiae. Yeast 11:1265–1274.

Schnell, U., F. Dijk, K. A. Sjollema and B. N. G. Giepmans, 2012 Immunolabeling artifacts and the need for live-cell imaging. Nature Methods 9:152–158.

Schwartz, M. A., and H. D. Madhani, 2004 Principles of MAP kinase signaling specificity in Saccharomyces cerevisiae. Annu Rev Genet 38:725–748.

Seger, R., D. Seger, A. A. Reszka, E. S. Munar, H. Eldar-Finkelman et al., 1994 Overexpression of mitogen-activated protein kinase kinase (MAPKK) and its mutants in NIH 3T3 cells. Evidence that MAPKK involvement in cellular proliferation is regulated by phosphorylation of serine residues in its kinase subdomains VII and VIII. J Biol Chem 269:25699–25709.

Shapiro, P. S., E. Vaisberg, A. J. Hunt, N. S. Tolwinski, A. M. Whalen et al., 1998 Activation of the MKK/ERK pathway during somatic cell mitosis: direct interactions of active ERK with kinetochores and regulation of the mitotic 3F3/2 phosphoantigen. J Cell Biol 142:1533–1545.

Sharma, V., 2018 ImageJ plugin HyperStackReg V5.6 (Version v5.6), pp. Zenodo.

Shaul, Y. D., and R. Seger, 2006 ERK1c regulates Golgi fragmentation during mitosis. J Cell Biol 172:885–897.

Shaul, Y. D., and R. Seger, 2007 The MEK/ERK cascade: from signaling specificity to diverse functions. Biochim Biophys Acta 1773:1213–1226.

Shaulian, E., and M. Karin, 2001 AP-1 in cell proliferation and survival. Oncogene 20:2390–2400.

Sidorova, J., and L. Breeden, 1993 Analysis of the SWI4/SWI6 protein complex, which directs G1/S-specific transcription in Saccharomyces cerevisiae. Mol Cell Biol 13:1069–1077.

Sikorski, R. S., and P. Hieter, 1989 A system of shuttle vectors and yeast host strains designed for efficient manipulation of DNA in Saccharomyces cerevisiae. Genetics 122:19–27.

Slater, M. L., 1973 Effect of reversible inhibition of deoxyribonucleic acid synthesis on the yeast cell cycle. J Bacteriol 113:263–270.

Slater, M. L., 1974 Recovery of yeast from transient inhibition of DNA synthesis. Nature 247:275–276.

Spellman, P. T., G. Sherlock, M. Q. Zhang, V. R. Iyer, K. Anders et al., 1998 Comprehensive identification of cell cycle-regulated genes of the yeast Saccharomyces cerevisiae by microarray hybridization. Mol Biol Cell 9:3273–3297.

Stevenson, B. J., N. Rhodes, B. Errede and G. F. Sprague, Jr., 1992 Constitutive mutants of the protein kinase STE11 activate the yeast pheromone response pathway in the absence of the G protein. Genes Dev 6:1293–1304.

Strickfaden, S. C., M. J. Winters, G. Ben-Ari, R. E. Lamson, M. Tyers et al., 2007 A mechanism for cell-cycle regulation of MAP kinase signaling in a yeast differentiation pathway. Cell 128:519–531.

Taheri, N., T. Köhler, G. H. Braus and H. U. Mösch, 2000 Asymmetrically localized Bud8p and Bud9p proteins control yeast cell polarity and development. Embo j 19:6686–6696.

Tanaka, K., K. Tatebayashi, A. Nishimura, K. Yamamoto, H. Y. Yang et al., 2014 Yeast osmosensors Hkr1 and Msb2 activate the Hog1 MAPK cascade by different mechanisms. Sci Signal 7:ra21.

Tatebayashi, K., K. Tanaka, H. Y. Yang, K. Yamamoto, Y. Matsushita et al., 2007 Transmembrane mucins Hkr1 and Msb2 are putative osmosensors in the SHO1 branch of yeast HOG pathway. Embo j 26:3521–3533.

Tatebayashi, K., K. Yamamoto, M. Nagoya, T. Takayama, A. Nishimura et al., 2015 Osmosensing and scaffolding functions of the oligomeric four-transmembrane domain osmosensor Sho1. Nat Commun 6:6975.

Tatebayashi, K., K. Yamamoto, K. Tanaka, T. Tomida, T. Maruoka et al., 2006 Adaptor functions of Cdc42, Ste50, and Sho1 in the yeast osmoregulatory HOG MAPK pathway. Embo j 25:3033–3044.

Thévenaz, P., U. E. Ruttimann and M. Unser, 1998 A pyramid approach to subpixel registration based on intensity. IEEE Trans Image Process 7:27–41.

Vadaie, N., H. Dionne, D. S. Akajagbor, S. R. Nickerson, D. J. Krysan et al., 2008 Cleavage of the signaling mucin Msb2 by the aspartyl protease Yps1 is required for MAPK activation in yeast. J Cell Biol 181:1073–1081.

Vallen, E. A., J. Caviston and E. Bi, 2000 Roles of Hof1p, Bni1p, Bnr1p, and myo1p in cytokinesis in Saccharomyces cerevisiae. Mol Biol Cell 11:593–611.

van der Felden, J., S. Weisser, S. Brückner, P. Lenz and H.-U. Mösch, 2014 The Transcription Factors Tec1 and Ste12 Interact with Coregulators Msa1 and Msa2 To Activate Adhesion and Multicellular Development. Molecular and Cellular Biology 34:2283.

Vandermeulen, M. D., and P. J. Cullen, 2020 New Aspects of Invasive Growth Regulation Identified by Functional Profiling of MAPK Pathway Targets in Saccharomyces cerevisiae. Genetics.

Vinod, P. K., N. Sengupta, P. J. Bhat and K. V. Venkatesh, 2008 Integration of global signaling pathways, cAMP-PKA, MAPK and TOR in the regulation of FLO11. PLoS One 3:e1663.

Waltermann, C., M. Floettmann and E. Klipp, 2010 G1 and G2 arrests in response to osmotic shock are robust properties of the budding yeast cell cycle. Genome Inform 24:204–217.

Wasch, R., and F. R. Cross, 2002 APC-dependent proteolysis of the mitotic cyclin Clb2 is essential for mitotic exit. Nature 418:556–562.

White, M. A., L. Riles and B. A. Cohen, 2009 A systematic screen for transcriptional regulators of the yeast cell cycle. Genetics 181:435–446.

Wilson, W. A., S. A. Hawley and D. G. Hardie, 1996 Glucose repression/derepression in budding yeast: SNF1 protein kinase is activated by phosphorylation under derepressing conditions, and this correlates with a high AMP:ATP ratio. Curr Biol 6:1426–1434.

Wittenberg, C., and S. I. Reed, 2005 Cell cycle-dependent transcription in yeast: promoters, transcription factors, and transcriptomes. Oncogene 24:2746–2755.

Wright, J. H., E. Munar, D. R. Jameson, P. R. Andreassen, R. L. Margolis et al., 1999 Mitogen-activated protein kinase kinase activity is required for the G(2)/M transition of the cell cycle in mammalian fibroblasts. Proc Natl Acad Sci U S A 96:11335–11340.

Yamamoto, K., K. Tatebayashi and H. Saito, 2016 Binding of the Extracellular Eight-Cysteine Motif of Opy2 to the Putative Osmosensor Msb2 Is Essential for Activation of the Yeast High-Osmolarity Glycerol Pathway. Mol Cell Biol 36:475–487.

Yamamoto, K., K. Tatebayashi, K. Tanaka and H. Saito, 2010 Dynamic control of yeast MAP kinase network by induced association and dissociation between the Ste50 scaffold and the Opy2 membrane anchor. Mol Cell 40:87–98.

Yang, H. Y., K. Tatebayashi, K. Yamamoto and H. Saito, 2009 Glycosylation defects activate filamentous growth Kss1 MAPK and inhibit osmoregulatory Hog1 MAPK. Embo j 28:1380–1391.

Yofe, I., U. Weill, M. Meurer, S. Chuartzman, E. Zalckvar et al., 2016 One library to make them all: streamlining the creation of yeast libraries via a SWAp-Tag strategy. Nat Methods 13:371–378.

Yoon, S., and R. Seger, 2006 The extracellular signal-regulated kinase: multiple substrates regulate diverse cellular functions. Growth Factors 24:21–44.

Zarzov, P., C. Mazzoni and C. Mann, 1996 The SLT2(MPK1) MAP kinase is activated during periods of polarized cell growth in yeast. Embo j 15:83–91.

Zehorai, E., Z. Yao, A. Plotnikov and R. Seger, 2010 The subcellular localization of MEK and ERK--a novel nuclear translocation signal (NTS) paves a way to the nucleus. Mol Cell Endocrinol 314:213–220.

